# The viscoelastic properties of *Nicotiana tabacum* BY-2 suspension cell lines adapted to high osmolarity

**DOI:** 10.1101/2024.04.29.591580

**Authors:** Tomasz Skrzypczak, Mikołaj Pochylski, Magdalena Rapp, Przemysław Wojtaszek, Anna Kasprowicz-Maluśki

## Abstract

To survive and grow, plant cells must regulate the properties of their cellular microenvironment in response to ever changing external factors. How the biomechanical balance across the cell’s internal structures is established and maintained during environmental variations remains a nurturing question. To provide insight into this issue we used two micro-mechanical imaging techniques, namely Brillouin light scattering and BODIPY-based molecular rotors Fluorescence Lifetime Imaging, to study *Nicotiana tabacum* suspension BY-2 cells long-term adapted to high concentrations of NaCl and mannitol. We discuss our results in terms of molecular crowding in cytoplasm and vacuoles, as well as tension in plasma membrane. The viscoelastic behavior was elucidated relative to cells external environments revealing the difference between the responses of cytoplasm and vacuole in the adapted cells. To understand how sudden changes in osmolarity affect cellular mechanics, the response of control and already adapted cells to further short-term osmotic stimulus was also examined. The applied correlative approach provides evidence that adaptation to hyperosmotic stress leads to different ratios of protoplast and environmental qualities that help to maintain cell integrity. Presented results demonstrate that the viscoelastic properties of protoplasts are an element of plant cells adaptation to high osmolarity.

## Introduction

Understanding the interactions between the cellular microenvironment and the external environment remains a significant challenge in cell biology. One fundamental environmental factor with pleiotropic impacts is osmolarity, which can alter water activity and directly affect all cellular processes and biomechanics of a plants cell (Watson et al., 2023; Zonia & Munnik, 2007). Despite the vast information on the response to stress, it is still elusive how much changes in the properties of biophysical cells originate from the direct impact of an environment (i.e. passively imposed), and from the activity of cells in response to stress. Hyperosmolarity, as well as desiccation and temperature fluctuations, could severely altered water activity in cells, what disturbs cell growth, and entire molecular machinery, which necessitates reactions to overcome dysfunctions (Romero-Perez et al., 2023; van Zelm et al., 2020; Watson et al., 2023; Zonia & Munnik, 2007). Recent studies have demonstrated that hyperosmolarity stress rapidly increases cellular viscosity and molecular crowding in neuroblastoma cells (Kitamura et al., 2023). Fundamentally, hyperosmotic stress explicitly leads to increased molecular crowding (MC), due to water efflux, which dramatically alteres conditions, that are crucial for cellular molecular machinery (Romero-Perez et al., 2023). Moreover, a quick water efflux from a cell could induce temporal or irreversible plasmolysis. During osmotic and salt stress at first signalling pathways are activated at first, leading to a quick stress response; in time, cells adjust their physiology, what include growth quiescence and recovery (reviewed in Julkowska & Testerink, 2015; Zhu, 2002). In the case of salt stress, an excessive concentration of sodium is toxic, also by altering cation homeostasis (K^+^/Na^+^) and plasma membrane potential (Li & Yang, 2023). Additionally, also a high concentration of chloride anion is toxic and disturb anions homeostasis and consequently organellar physiology (Geilfus, 2018).

With prolonged increased salinity, plants can develop adaptations at the cellular level, including increased biosynthesis of osmolytes, cell wall remodelling, and intense export of Na^+^ to vacuoles (Flowers & Colmer, 2015; Li & Yang, 2023; van Zelm et al., 2020; Zhao et al., 2020). The viscosity, which can also change due to hyperosmolarity, was shown to directly impact metabolism and enzymatic reactions (Gavish & Werber, 1979; Kitamura et al., 2023; Uribe & Sampedro, 2003). However, the viscoelastic properties of cells, although crucial for interactions between cells and environment, are much less studied than networks of signalling pathways and transcriptional response. It was shown, that in the mammalian cells, the cytoplasm behaves differently, like liquid or like a gel, for molecules of different size. Importantly, the cytoplasmic viscosities were found as similar across various cells lines (Kwapiszewska et al., 2020). The reason that the local viscosity and MC must be kept within a certain range is to clear the path for efficient diffusion (Alfano et al., 2023). In yeasts, it was shown that ‘viscoadaptation’ depends on the synthesis of glycogen and trehalose. Importantly, ‘viscoadaptation’ allows yeast cells to maintain efficient diffusion over a wide temperature range (Persson et al., 2020). In HeLa cells, changes in cell volume in response to osmotic shocks are followed by changes in membrane tension, and such volume-membrane tension dynamics are actively regulated by the cell (Roffay et al., 2021). Recently, it was shown, that fundamentally for the stress response, biocondensates made of Intrinsically Disordered Proteins (IPDs) counteract microenvironmental instabilities induced by temperature and osmolarity changes (Watson et al., 2023).

The turgor pressure and the bulk elastic modulus of a plant cell can be measured directly with a microcapillary invasive pressure probe (Tomos & Leigh, 1999). Less-invasive approach for measuring elastic and viscoelastic properties of plant cells are indentation methods, based on the idea of measuring the force required to displace the surface of a sample by a given distance. The resulting force versus indentation curves can then be used to calculate the stiffness of a sample. These include atomic force microscopy (AFM) (Milani et al., 2011; Peaucelle et al., 2011; Sampathkumar et al., 2014), cellular force microscopy (CFM) (Routier-Kierzkowska et al., 2012), microcompression (Wang, 2004; Wang et al., 2006), and other single-point indentation systems (Hayot et al., 2012; Lintilhac et al., 2000).

In the last decade many techniques have been developed and utilised to measure the mechanical properties of living biological materials, which enable *in situ* measurements (Elsayad et al., 2016; Farokh Payam & Passian, 2024; Kitamura et al., 2023; Michels et al., 2020; Robinson et al., 2023; Seelbinder et al., 2024; Wu et al., 2018). Cell microenvironment-sensing dyes based on fluorescence lifetime (FLIM) signals have proven to be robust and independent from concentration, precision in revealing cellular characteristics, such as pH, intrinsically disordered proteins status, redox status, hormone concentrations (Colin et al., 2022; Cuevas-Velazquez et al., 2021; Kitamura et al., 2023; Koveal et al., 2020; Miller et al., 2020; Steinbeck et al., 2020; van der Linden et al., 2021). Importantly for plant biomechanics research, BODIPY-based fluorescent rotors were shown as efficient tools in mapping viscosity with subcellular precision (Michels et al., 2020).

Brillouin Light Scattering (BLS) was introduced as an all-optical contactless mechanical probe with diffraction-limited spatial resolution. This optical technique rely on the scattering of light on thermally generated acoustic waves, ever-existing in any material. These waves acts as a mechanical stimuli, making material to effectively constantly deform itself, which allow to extract viscoelastic information from spectroscopic data. Being fully non-invasive, label-free and contactless, this technique can deliver mechanical information when physical access to the region of interest cannot be gained, such as in a cell or multicellular object (Elsayad et al., 2016; Gouveia et al., 2019; Margueritat et al., 2019; Roberts et al., 2021; Wisniewski et al., 2020; Bevilacqua et al., 2023; Elsayad et al., 2016). Recently, the method was also used to study the stiffness of the cell walls of plants at the cellular level in roots of Arabidopsis thaliana (Bacete et al., 2022; McKenna et al., 2019). Although the application of Brillouin microscopy might allow the determination of the elastic factor of a solid body, in soft matter/hydrated materials, such as cells or polymer solutions, the water content might be more reliably measured (P. J. Wu et al., 2018). BLS was also revealed as an efficient tool for investigating molecular crowd dynamics in multicellular spheroids in response to hyperosmolarity (Yan et al., 2021) and recently to study mechanical anisotropy in living matter (Keshmiri et al., 2024).

Despite growing number of studies devoted to viscoelastic behavior of cells, this property is still far less understood than networks of signaling pathways and transcriptional responses. In addition, the role of the surrounding medium (environment) on stress-induced changes in the cell’s biomechanics remains unclear. It is still elusive to which extent viscoelastic state of cell is directly related to mechanics of its environment or follows from the activity of cell during its long term adaptive response to the stress imposed by an environment.

The tobacco (*Nicotiana tabacum*) BY-2 suspension root cell line is a well-established model in cell biology and biotechnology research (Hoque et al., 2007; Jelínková et al., 2015; Khatun et al., 2020; Kurusu et al., 2012; Olmos et al., 2017; Srba et al., 2016). The use of BY-2 allows for investigation of cells derived from relatively homogenous population, that live in homogenous environments (culture media). Here, we utilised BY-2 cell lines that were adapted to high concentrations of NaCl and mannitol. Gradual adaptations were performed as far as the highest tolerated concentrations were obtained, i.e. 190 mM NaCl for NaCl adapted BY-2 (BY-2:NaCl), and 450 mM mannitol for mannitol adapted BY-2 (BY-2:Mann). The adapted cell lines obtained have survived for multiple generations. The adapted cell lines presented significantly different morphologies and transcriptomes from the control cell lines and from each other as well (Skrzypczak et al., 2024).

In trying to address mechanisms of osmoadaptation, we performed micro-viscoelastic imaging on BY-2 suspension root cell line under various osmotic conditions. The utilisation of BODIPY(BDP)-based molecular rotors and BLS enabled us to reveal viscoelastic BY-2 properties with subcellular precision and by non-destructive methods. We utilised cells exposed to high concentrations of NaCl or mannitol in the short-term (‘stress’) or long-term conditions (‘adaptation’) to establish a useful model to elucidate cellular responses to hyperosmolarity.

## Materials and methods

### BY-2 cultures

Tobacco BY-2 suspension-cultured cells were maintained in liquid medium, pH 5.0, containing Murashige and Skoog salts (Murashige & Skoog 1962) and (per liter): 30 g sucrose, 100 mg inositol, 3 mg 2,4-D, 370 mg KH_2_PO_4_ and 1 mg thiamine-HCl (Nagata *et al*. 1992). Cells were subcultured weekly at 10 ml of culture per 70 ml of fresh medium in 300-mL Erlenmeyer flasks. Cultures were incubated at 21°C on a gyratory shaker at 120 rpm with a 2.5-cm displacement. During 18 months, the cells were gradually adapted to grow in a media supplemented with either 450 mM mannitol (BY-2:Mann) or 190 mM NaCl (BY-2:NaCl). The cultures were transferred to fresh media containing increasing concentration of mannitol or NaCl in sequential manner, starting with 50 mM mannitol and 20 mM NaCl (osmotic agents concentrations were increased every 2 months), but were otherwise handled identically to the BY-2:Control cells. Immediately following transfer, the decrease intensity of cell divisions and decrease in the fresh weight of cells in culture were noted. However, after 4-8 subsequent passages, the adapted cells exhibited the same growth properties as control ones. This was usually considered appropriate the time point for the subsequent increase in the concentration of osmotic agent in culture media. Final concentrations of mannitol (450 mM) and NaCl (190 mM) were the highest that were not lethal to BY-2 cells. The osmolarities of fresh media were measured with an osmometer. The highest media osmolarity was measured in the high mannitol adapted line (BY-2:Mann; ∼750 mOsm/kg), than in the high NaCl adapted line (BY-2:NaCl; ∼550 mOsm/kg). The lowest osmolarity was measured for the standard BY-2 medium used to culture BY-2:Control; ∼220 mOsm/kg.

All the results described herein are from cultures maintained in the presence of final concentrations of osmotic agents for at least 15 years, and all the sampling and stress treatments were done on the third day (logarithmic growth phase) after transfer to fresh medium. For stress experiments, NaCl and mannitol were added to medium derived by centrifugation from the same day of culture. This allowed for minimising the effects of media compositions different from those of stress agents. To improve synchronization, cultures were kept for 2 weeks without subculture, followed by dilution of 5 ml of old culture in 70 ml of fresh medium. Undifferentiated and dividing cells were observed 2 days after that subculture, while differentiated non-dividing cells were observed after 10 days of culture.

### Fluorescent BODIPY rotors synthesis

BODIPY-based (BDP) molecular rotors were synthesised according to a protocol described previously (Michels et al., 2020). Procedure and NMR data results of synthesised rotors are available in Supplement: <u>SI_BDP_synthesis.pdf</u>

### BODIPY rotors imaging and data analysis

Cellular microviscosity was measured with BDP-based molecular rotor fluorescence probes in the middle days of the culture cycles. The cells were incubated with PEG-BDP for 1 h and with N^+^-BDP for 3 h, and then washed twice with their respective culture media derived from a BY-2 culture. The PicoHarp300-Dual Channel SPAD system (PicoQuant) in combination with a Nikon A1Rsi microscope armed with the 20x dry or 60x /1.20 water immersion objective was used. The Picosecond Pulsed Diode Laser LDH-D-C-485 and Supercontinuum Tunable White Light Laser (488 nm) were used for generation of 100 ps excitation pulses at a repetition of 40 MHz. The images were of frame size of 256 × 256 pixels and were collected with an average count rate of around 10^5^ photons per second for each apprx. 30-60 s, so that maximum photons per pixel reached 3 000 counts. FLIM data were collected from more than 30 cells. Regions of Interest (ROI) were selected to mark properly subcellular compartments. Furthermore, also cells-free areas, but the surrounding area of the cells was used to obtain a real lifetime signal of the environment of BY-2 cells (media). The fluorescence decay curves were fitted in each pixel with a single-component exponential decay, and the lifetime mean and SD were calculated. The lifetime signals distributions were assigned to 0.1 ns wide bins, then events occurrence scaled in (0, 1) range and plotted with matplotlib as histograms.

### Brillouin light scattering

In the Brillouin light scattering (BLS) experiment the incident laser beam from laser (Spectra Physics, Excelsior, λ=532nm) was delivered to custom-built microscope of the usual design (Fig.1a). The beam of vertical polarisation, reflected by the polarising beam-splitter, was passed through λ/4 to produce incident beam of circular polarisation. The beam was focused on the sample with a 20x microscope objective (Zeiss LD Achroplan Korr; NA=0.4). A low numerical aperture objective was used to reduce the spectral broadening effect (Antonacci et al., 2013) with the consequence of increasing the volume of scattering (6×20 um). The sample was moved in the XY plane with a motorised translation stage (Märzhäuser Wetzlar GmbH & Co. KG). The power measured on the sample was 10 mW. The light backscattered from the sample was collected by the same objective and after restoring the vertical polarization (by passing through λ/4) was transmitted by beam-splitter towards λ/2 used to rotate polarization plane of scattered light before focussing on the entrance pinhole of the Brillouin spectrometer. For Brillouin spectrum measurement, Tandem Fabry-Perot Interferometer (TFPI) (TFP2, The Table Stable Ltd.) whose spectral range (FSR) was set to 15 GHz (10 mm mirror spacing).

**Fig. 1.**
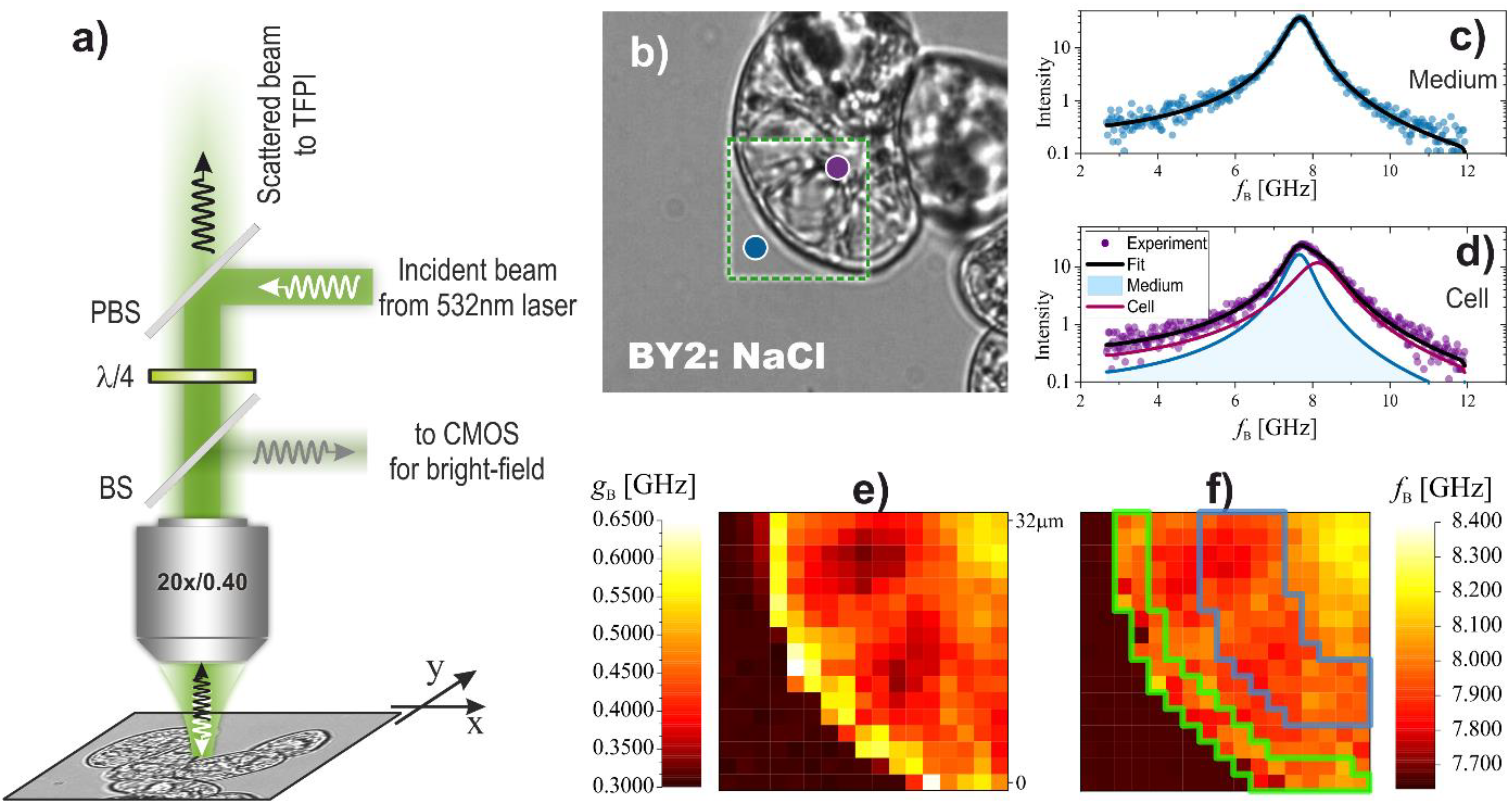
Brillouin imaging of a cell. a) Schematic of the experimental setup for Brillouin micro-spectroscopy. BS, PBS, λ/4 and λ/2 stand, respectively, for removable beam splitter (for bright-field imaging), polarising beam splitter, quarter- and half-wave plates. b) Bright-field (BF) image of the cell. The green rectangle indicates the area chosen for Brillouin imaging. The blue and violet circles show the positions of the buffer and interior of the cell, respectively. c) Brillouin spectrum (anti-Stokes side) collected for buffer (Blue circle in b)). Notice that a single Brillouin peak is observed. d) Brillouin spectrum acquired inside the cell (violate circle in b)). Notice that two peaks are necessary to reproduce the spectrum. The first peak is assumed to come from scattering from a buffer i.e. medium (blue line). The second is a signal from the cell interior (dark red line). During spectrum fitting, the Brillouin peak parameters of the first peak were fixed to the values found for buffer. The Brillouin line shift and line width of the second peak (corresponding the cell) were mapped. e) Map of the Brillouin linewidth. f) Map of the Brillouin lines shift. Green line border region identified in BF image as cytoplasm dominating. Blue line border region identified in BF as vacuole dominating. The size of the maps is 32um.

The sample was sandwiched between two microscope slides. The starting position for XY scanning was chosen after inspection of the sample using bright-field imaging with the same microscope. At given XY position, recorded spectrum was average of 15 spectral sweeps giving ∼10 s for single spectrum acquisition. Spectra were collected for a 16×16 grid spaced by 2 um. The total time to collect a single map was ∼45 min. Each map was measured for a fresh sample.

In the investigation of cell mechanics, cells from every BY-2 line were extensively mapped, with more than 8 cells analysed for each line. The mapping process involved capturing bright field images to select specific regions showing bulk buffer, cell wall, and cell interior visibility, as illustrated in Fig.1b.

Brillouin spectra analysis was conducted using single-phase or two-phase model (<u>SI_BLS.pdf</u>; Eq.S1), depending on whether the mapped area corresponded, respectively, to the bulk medium (Fig.1c),or cell interior (Fig.1d). The spatial distribution of the Brillouin line shift, *f*_B_ (informing about longitudinal elasticity), and the Brillouin line width, *g*_B_ (quantifying longitudinal viscosity), was visually represented in the form of maps (Fig.1e,f), allowing for a comprehensive understanding of their variations across the cell. To enhance the analytical depth, manual selection of pixels was performed in each map, identifying three distinct regions: 1) medium (cell exterior/environment), 2) vacuole (interior of the cell, excluding nuclear region), and 3) cytoplasm (a protoplast in the vicinity of the cell wall).

Further insights into the viscoelastic properties of the cell and its environment were obtained through the production of histograms for *f*_B_ and *g*_B_ values. The parameters of the Brillouin line, such as shift and width, were examined for the three identified regions, providing a statistical overview of cellular mechanics and its surrounding medium. This multifaceted approach not only enabled a detailed analysis of individual cells, but also contributed to a broader understanding of the viscoelastic characteristics across different BY-2 lines.

## Results

### Adapted BY-2 lines exhibited unique properties of cytoplasmic molecular crowd

BDP-based cytoplasmic and plasma membrane (PM) based molecular rotors were used to obtain information on the viscosity of BY-2 subcellular compartments. The signals derived from PEG-BDP in the cytoplasm could be interpreted as an indicator of local molecular crowd (MC), while N^+^-BDP works as a sensor of plasma membrane (PM) fluidity (Michels et al., 2020). In the media, that is a liquid environment, PEG-BDP provided data that correspond to viscosity (Michels et al., 2020), so the cytoplasmic molecular crowd was presented relative to environmental viscosity.

PEG-BDP efficiently stained the cytoplasm of BY-2 cells and did not penetrate into vacuoles, but clearly diffused also to the nucleus (Fig. 2A). ROIs were used to isolate signals from the cytoplasm and to exclude denser nuclear areas. Lifetime was also measured in the medium area in cells vicinity (∼50-100 um from cells) to respect the impact of the real environment on the cellular microenvironment. This allowed the calculation relative of the differences between cells and their environment (Δ; <u>SI_BDPimage.pdf</u>). The lifetimes measured in the surrounding media were smaller than those measured inside the cells, and the lifetime distributions of the media were clearly distinguishable from those of the cells (Table 1).

**Table 1.**
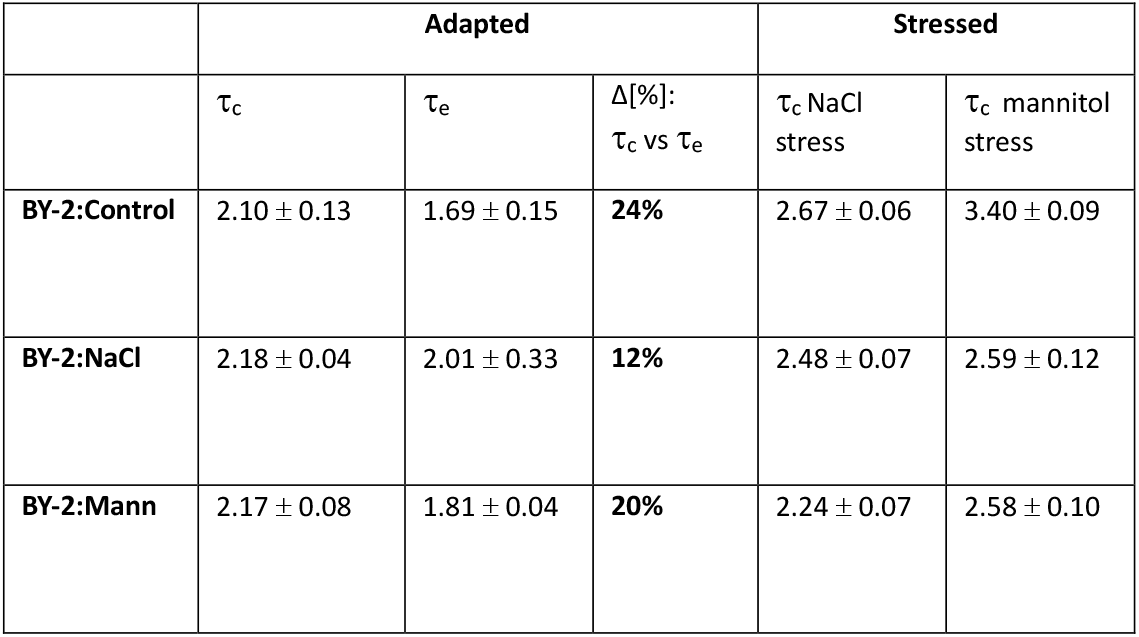
Means and standard deviations (±) of lifetime PEG-BDP values derived from cytoplasm (τ_c_) of BY-2 cells and medium (τ_e_) in cells vicinity. The relative difference between cells and media was calculated and presented as Δ[%]. All values in ns.

**Fig. 2.**
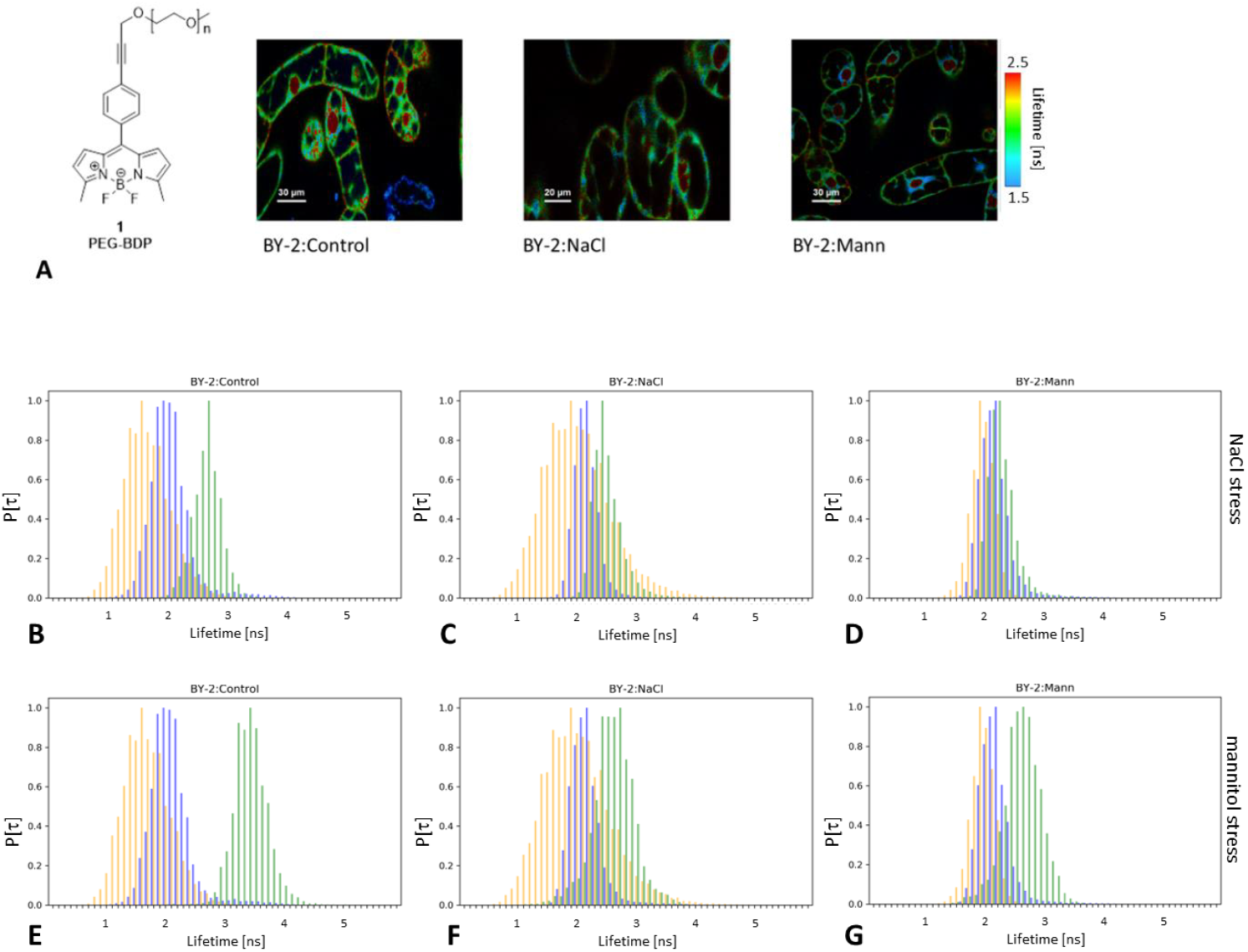
A) PEG-BDP formula and images of PEG-BDP stained BY-2 lines. Lifetime BY-2 images are on a false colour scale that represents the mean fluorescence lifetime for each pixel, expressed in nanoseconds. Nuclear regions with significantly higher lifetime values were not considered to determine the cytoplasmic lifetime and molecular crowd; B-G) Normalised distributions of the PEG-BDP lifetime derived from the cytoplasm (τ_c_) of the BY-2 lines. The cytoplasmic signal (blue) is plotted together with the medium signal (orange, τ_e_) derived from cell vicinities and the cytoplasmic signal for the stressed cells (green). (A-C) 200 mM NaCl stress; (D-F) 450 mM mannitol stress. A,D) BY-2:Control; B,E) BY-2:NaCl; C,F) BY-2:Mann

The interior of the cells was expected to be denser and more viscous than the surrounding media. Indeed, it was already confirmed in BY-2:Control, in which lifetime of PEG-BDP was higher in the cytoplasm (τ_c_∼2.1 ns), than in the media (τ_e_∼1.69 ns). This translated to the relative difference (Δ) between the cytoplasm and the media around 24% (Table 1).

The adaptation to media of higher osmolarity requires that cells counteract the change and try to preserve intracellular conditions that allow for efficient diffusion and balance of water flow between a cell and an environment. Both adapted BY-2 lines displayed only slightly different PEG-BDP lifetime signals distributions, than did BY-2:Control. The distributions of lifetime values were slightly shifted toward higher lifetimes; +0.07-0.08 ns (Table 1). On the other hand, the relative to medium lifetime values (Δ) are much smaller in the both adapted BY-2 lines than in BY-2:Control. The lifetime signal distributions of the adapted cells were much closer to the media signals (Fig. 2). The relative lifetime (Δ) of PEG-BDP in the cytoplasm was the highest in BY-2:Control (24%), followed by BY-2:Mann (20%) and BY-2:NaCl (12%) (Table 1). This effect was more manifested in fluorescence lifetime distributions in media (τ_e_), than in cytoplasm (τ_c_) (Fig. 2).

Despite the fact that cells had been living in environment of higher osmolarity for many generations, the cytoplasmic molecular crowding remained almost stable. The smaller Δ in adapted BY-2:NaCl cells would have implied a cytoplasm of more similar density in comparison to environment, than in the case of BY-2:Control.

Interestingly, although, the τ_e_ was the highest for BY-2:NaCl, it did not correspond to the measured media osmolarities, which was the highest for the BY-2:Mann media (Table 1). Because of the varied molecules’ size and the electric charges, mannitol and NaCl might affect the osmolarity and viscosity of the medium differently. In two different environments with high osmolarity only slightly altered cytoplasmic MC were observed.

Then, the osmotic stress response of BY-2 lines was determined. Increasing the NaCl concentration (+200 mM) led to the moderate level of plasmolysis within an hour of all BY-2 lines. Such salt stress resulted in significantly altered PEG-BDP lifetime distributions in BY-2:Control in a timeline below 30 min (Table 1; Fig. 2A-C; <u>SI_BDPimage.pdf</u>: SI_Fig. 2). The τ_c_ went significantly higher (τ_c_ + 0.57 ns). This corresponds to an increase in MC in response to increased media osmolarity. The water efflux from the cells was induced. Similar, but more prominent trends (τ_c_ + 1.3 ns), were observed after an application of 450 mM mannitol (Table 1). The addition of extra mannitol led to severe plasmolysis within all lines, so massive water leakage from cells also had occurred, what rapidly increased crowding.

In both adapted lines increase in relative lifetime (Δ), as well as τ_c_ increments were at least twice much smaller in adapted lines, than in BY-2:Control (Table 1). MC increased the most in BY-2:Control, which lived in media of the lowest osmolarity and viscosity. Respectively, the changes were the smallest after stresses in BY-2:Mann, which were living in the media of the highest osmolarity (Fig. 2; Table 1). These smaller MC changes are consistent with fact, that BY-2:NaCl and BY-2:Mann experienced relatively smaller increase in media osmolarity. Thus, we concluded, that after adaptation BY-2 maintained their basic properties of cytoplasm. However, MC changed less during response to osmotic stress.

The preservation of the cytoplasmic microenvironment within some range seems to be required for cell survival and adaptation to high osmolarity. Furthermore, the cytoplasmic microenvironment after adaptation appears to make BY-2 cells less prone to osmotic stress-induced cytoplasm condensing.

### Plasma membrane tension is similar within BY-2 cell lines

N^+^-BDP rotors were shown to be an efficient tool for determining changes in plasma membrane tension (PM) (Michels et al., 2020). In BY-2 heterogeneity of the lifetime within PM and intracellular membranes was observed (Fig. 3A). Increased N^+^-BDP lifetime suggests decreased membrane tension, what occurs during plasmolysis.

**Fig. 3.**
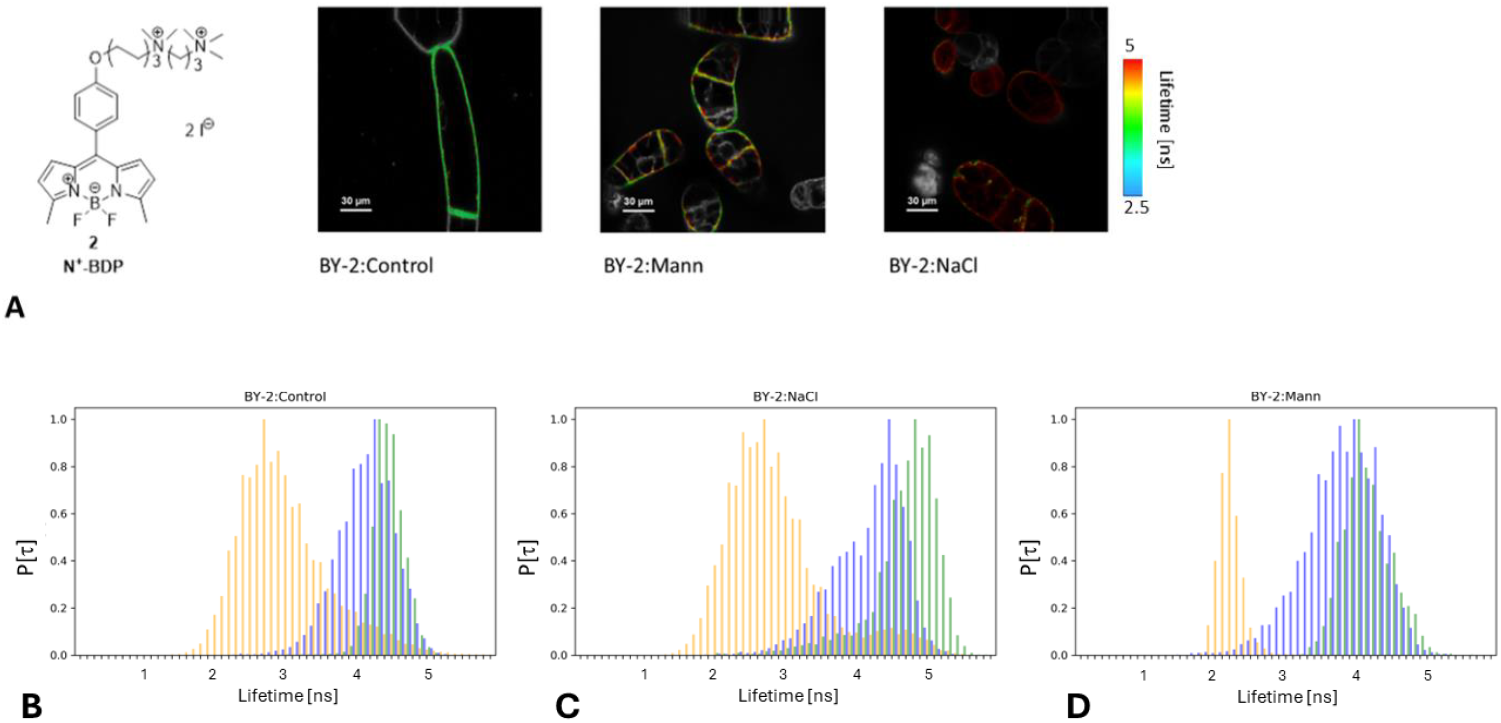
A) N_^+^_-BDP formula and N_^+^_-BDP stained BY-2 lines. The images are on false colour scale, representing the mean fluorescence lifetime for each pixel, expressed in nanoseconds. B-D) Normalised lifetime distributions of the N_^+^_-BDP signal derived from the plasma membrane (τ_m_) of BY-2 lines (blue), which were subjected to additional NaCl stress (green). The plasma membrane signal is plotted on the background of the media signal derived from the vicinity of the cells (orange, τ_me_). A) BY-2:Control; B) BY-2:NaCl; C) BY-2:Mann

The mean N^+^-BDP lifetime in PM of BY-2:Control was 4.16 ns. The PM lifetime distribution and the lifetime mean value (4.21 ns) were slightly higher in BY-2:NaCl than in BY-2:Control. Surprisingly, BY-2:Mann exhibited a significantly lower τ_m_ (3.82 ns), than BY-2:Control (Table 2; <u>SI_BDPimage.pdf</u>). This would suggest an increase of the PM tension in the BY-2:Mann. However, also, osmotic stress invoked by 450 mM mannitol led to decreased τ_m_. This would indicated greater PM fluidity and increased tension, as it would be expected in the case of hypoosmolarity, but not hyperosmolarity (Table 2). It is counterintuitive: a longer lifetime and corresponding lower tension of the wrinkled plasma membrane were expected in the results of plasmolysis, which was observed in the case of BY-2:Control stressed with mannitol. Our results suggested that high concentrations of mannitol disturbed the behaviour of N^+^-BDP. No such effect was observed for an increased NaCl concentration.

**Table 2.**
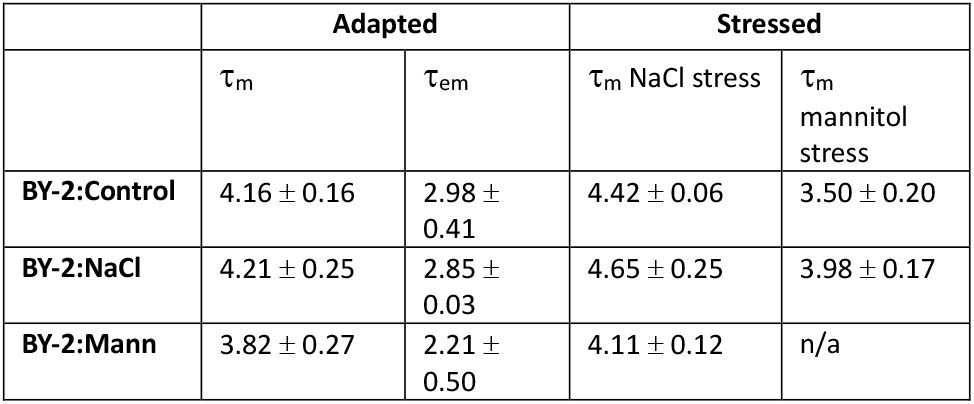
Mean values and standard deviation (±) values of N^+^-BDP lifetime values [ns] derived from plasma membrane (τ_m_) of BY-2 cells and medium (τ_em_) in cells vicinity. All values in ns.

|  | Adapted |  | Stressed |  |
| --- | --- | --- | --- | --- |
| | $\tau_m$ | $\tau_{em}$ | $\tau_m$ NaCl stress | $\tau_m$ mannitol stress |
| BY-2:Control | $4.16 \pm 0.16$ | $2.98 \pm 0.41$ | $4.42 \pm 0.06$ | $3.50 \pm 0.20$ |
| BY-2:NaCl | $4.21 \pm 0.25$ | $2.85 \pm 0.03$ | $4.65 \pm 0.25$ | $3.98 \pm 0.17$ |
| BY-2:Mann | $3.82 \pm 0.27$ | $2.21 \pm 0.50$ | $4.11 \pm 0.12$ | n/a |

The τ_m_ in PM of BY-2:Control cells after 200 mM NaCl stress was higher, than in the default medium (τ_m_ + 0.26 ns). This implicates decreased PM tension that occurred in response to hyperosmolarity, that correspond to observed moderate level of plasmolysis. N^+^-BDP lifetime exhibited higher values also in the PM of BY-2:NaCl (τ_m_ + 0.44 ns) and BY-2:Mann (τ_m_ + 0.29 ns) in response to 200 mM NaCl stress (Table 2; Fig. 3B-D). Interestingly, BY-2:NaCl have the most altered N^+^-BDP signals from plasma membranes after salt stress. This suggests that PM tension changed the most in salt stressed BY-2:NaCl. The base level of the τ_m_ is slightly higher in BY-2:NaCl, also τ_m_ in NaCl stress response is greater in BY-2:NaCl. We could speculate, that such effect depends on altered PM lipids composition or direct impact of monovalent ions.

Our results suggest that through the adaptation process, the PM tension is maintained in BY-2:NaCl cells. Salt stress reduced PM tension in both BY-2:Control and the adapted ones, what correlates with plasmolysis and PM detachment from cell walls. However, the PM of BY-2:NaCl might be a bit more prone to decrease of tension during salt stress.

### Brillouin imaging revealed differences in viscoelasticity between BY-2:Control and adapted lines

The distributions of the Brillouin line shift, f_B,medium_, and linewidth, g_B,medium_, for pure BY-2 media are presented in Fig. 4. The distributions of the media were narrow. Brillouin line shift and the linewidth for each medium were significantly higher than those observed in bulk water. The lowest values of both Brillouin measurable were found for BY-2:Control medium, a bit higher values correspond to BY-2:NaCl medium and the highest were found for BY-2:Mann medium. This result reflects the gradual increase in concentration and type of solutes in prepared culture media (BY-2:Control-> BY-2:NaCl-> BY-2:Mann).

**Fig. 4.**
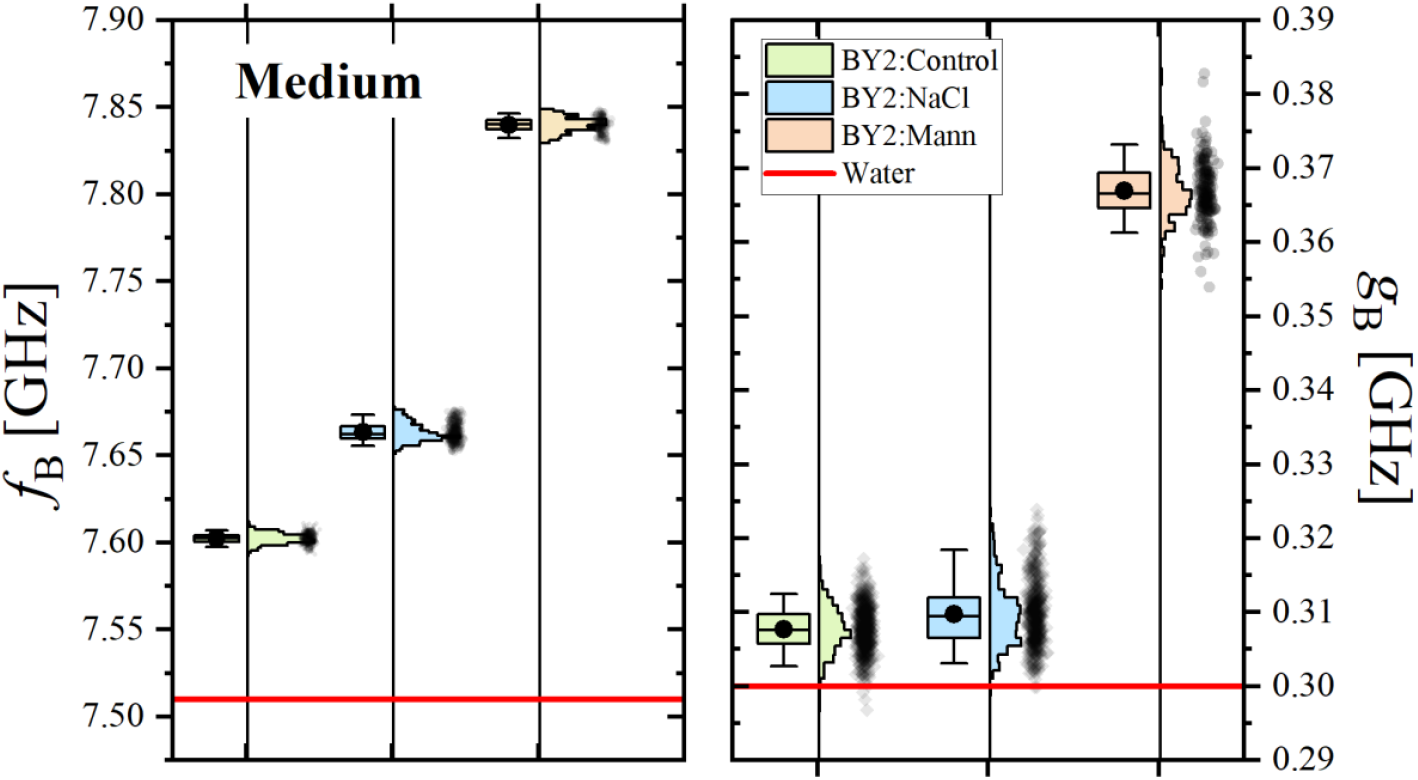
Distributions of Brillouin line shift, fB, and linewidth, gB, for pure media. Horizontal red lines indicate values for bulk water.

Distributions of Brillouin line shift and line width for vacuole and cytoplasm dominating areas (Fig. 1e) are presented on Fig. 5. The distributions of those values are much broader (than those for pure media) being the joint result of inhomogeneity of cells and variations in cells properties. The values of both Brillouin parameters are distinctly higher than those found in pure media buffers. This is the manifestation of a trivial fact that cells were much more concentrated than liquid media. Significantly, if the cell lives in a buffer characterised by higher Brillouin line-shift (or linewidth), its protoplast is also characterised by higher Brillouin measurable. The cells become more crowded, if it is exposed to a more concentrated environment of higher osmolarity. In result: water is relatively more excreted from the cell, or cells produce more molecules to increase molecular crowd. However, it is hard to say which of the above two has the highest contribution, since Brillouin is measurable depending on quantity (concentration) and quality (specific compounds dissolved) (Adichtchev et al., 2019; Yan et al., 2021).

**Fig. 5.**
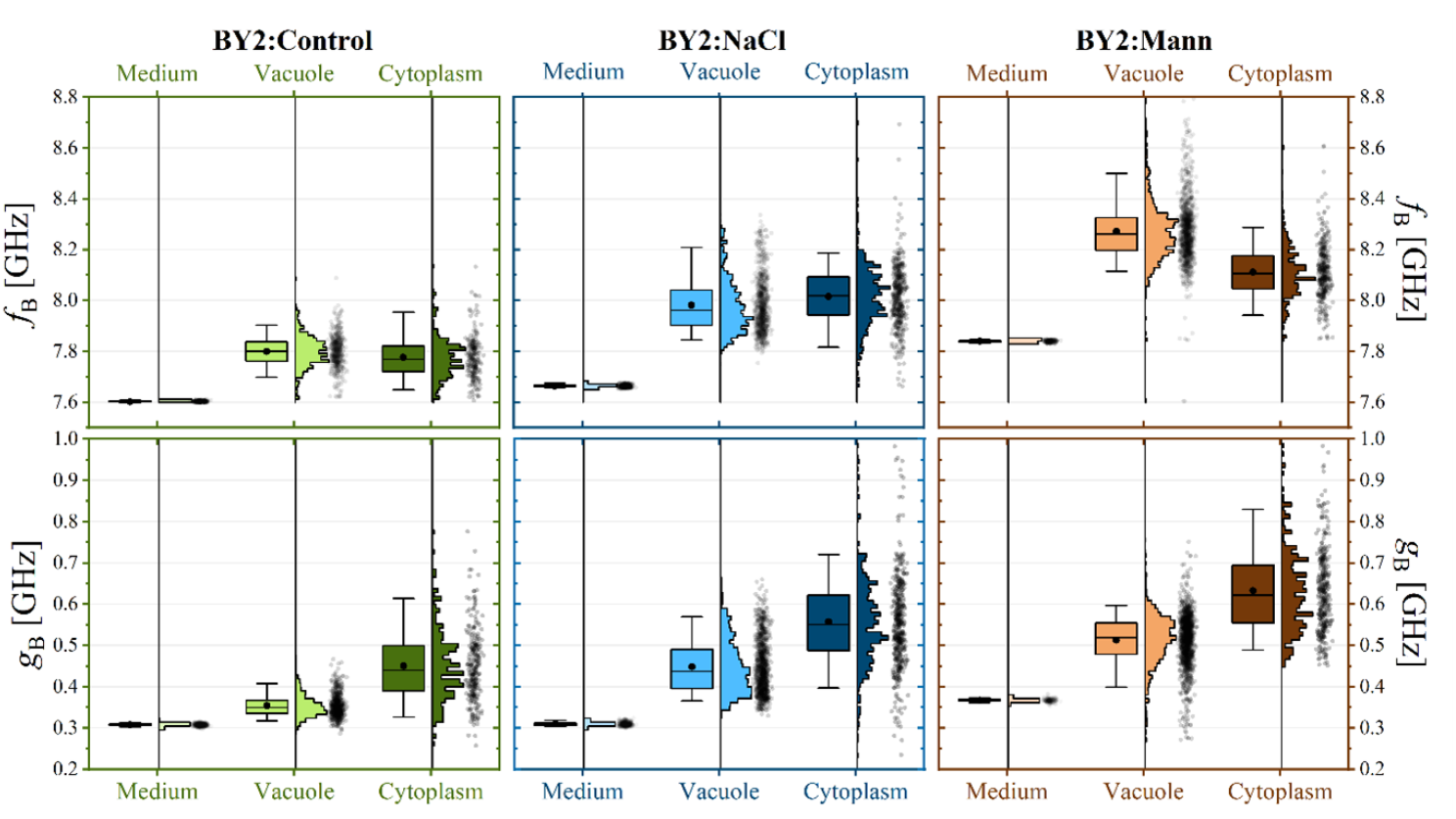
Distributions of the Brillouin line shift, f_B_,(upper panels) and Brillouin line width, g_B_, (lower panels) for pure media, cell vacuoles and cytoplasm for three cell lines of cells (BY-2:Control – green, BY-2:NaCl – blue, BY-2:Mann – brown).

To better visualise the effect of medium on cell mechanical properties, Brillouin results were presented in the form of the relative to the medium Brillouin shift (named ‘elastic contrast’ in (Bacete et al., 2021)), *υ*_B_, and the relative to the medium Brillouin line width (“viscous contrast”), *γ*_B_, according to (<u>SI_BLS.pdf</u>; x7). It should be emphasised that the definitions of mechanical contrasts (elastic and viscous) adopted in this study differ from the usual ones (Antonacci et al., 2020), where changes in Brillouin line shape parameters are given in relation to the values characterising pure bulk water. We decided to express the contrasts relative to the media, as these are the natural environments of the examined cells. Such defined mechanical contrasts can be interpreted in terms of change of internal cells composition being the result of cells adaptation to different environments.

The distributions of elastic and viscous contrasts are presented in Fig. 6.

**Fig. 6.**
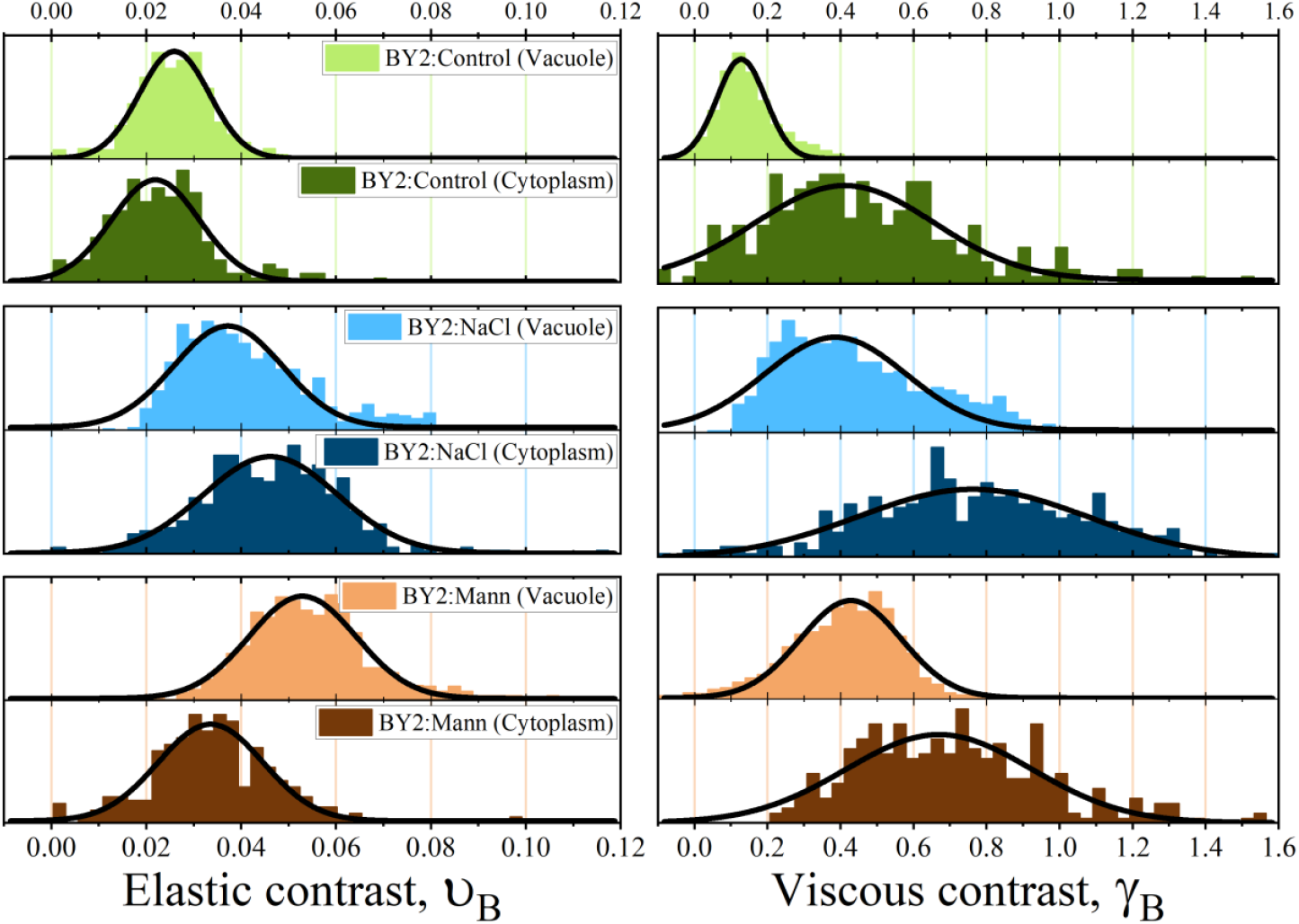
Distributions of relative Brillouin shift fB, and relative Brillouin line width, gB, for vacuoles dominating areas and cytoplasm dominating areas for three cell lines of cells (BY-2:Control – green, BY-2:NaCl – blue, BY-2:Mann – brown).

The BY-2 cell lines exhibited different Brillouin values *υ*_B_ and *γ*_B_. Few interesting observations can be made from the data presented in Fig. 6, and Table 3. The distributions of elastic and viscous contrasts for the vacuoles are the narrowest for BY-2:Control cells, suggesting that these are the most homogeneous cells on both the population scales and the single cell scales. The relative Brillouin shift (elastic contrast) found for the vacuoles of the BY-2:Control cells was about 2.5% higher than its medium. For BY-2:NaCl cells it was about 4% higher, whereas for cells living in high mannitol, BY-2:Mann, the mean value was 5%. If we recall that the Brillouin shift depends on the compressibility of the system (which changes with molecules concentration), and if we accept that vacuoles can be treated as an aqueous mixture, then we can interpret this result as an indication of molecular crowding. The effect was specific for a given culture medium, which is the highest for BY-2:Mann, followed by BY-2:NaCl, and the lowest for BY-2:Control. A qualitatively similar trend is observed for the viscous contrast (relative Brillouin width) obtained for the vacuoles. Here, the mean values of the distributions starting from 13% for BY-2:Control cells, followed by 39% for BY-2:NaCl cells. The highest values of 43% were again observed for BY-2:Mann.

**Table 3.** Elastic (υ_B_) and viscous (γ_B_) contrast values measured for different regions of the cells (cytoplasm and vacuole) belonging to three different lines. The values were obtained as a mean of the distribution (presented in Fig. 10 whereas the uncertainty (±) is the standard deviation.

| | $\nu_B$ [%] (Elastic contrast) | | $\gamma_B$ [%] (Viscous contrast) | |
| --- | --- | --- | --- | --- |
|  | Vacuole | Cytoplasm | Vacuole | Cytoplasm |
| BY-2:Control | $2.59 \pm 0.37$ | $2.19 \pm 0.93$ | $12.79 \pm 6.67$ | $41.15 \pm 24.55$ |
| BY-2:NaCl | $3.72 \pm 1.15$ | $4.62 \pm 1.42$ | $38.59 \pm 19.31$ | $76.19 \pm 31.34$ |
| BY-2:Mann | $5.29 \pm 1.13$ | $3.36 \pm 1.11$ | $42.83 \pm 13.83$ | $67.00 \pm 25.53$ |

The above analysis of the distributions of elastic and viscous contrasts suggests that long-term exposure of cells to environments of concentrated NaCl and mannitol made the vacuoles more crowded and viscous. This effect was the most pronounced in BY-2:Mann. The result of crowding was the reduction of the compressibility of the mixture and the increase in longitudinal viscosity. Brillouin values revealed variability in viscoelastic properties between cells adapted to various osmotic conditions. The elastic contrast values of vacuole are greater, than the elastic contrast values of cytoplasm. This effect was the most pronounced in BY-2:Mann. This might be consequences of the highest osmolarity to which were exposed BY-2:Mann, what might force the greatest mechanical rearrangements to maintain turgor and retain water.

Different results appeared when comparing the mechanical contrasts (both, elastic and viscous) recorded in cytoplasmic areas. Here, the highest values were found for BY-2:NaCl. Interestingly, BLS results showed that exposure to different culture media not only changed the overall composition of the cell, but also induced spatial inhomogeneity of this composition. This was partially confirmed by the observation that cells exposed to NaCl showed the broadest and clearly asymmetric distributions of mechanical contrasts (elastic and viscous) being skewed toward higher values. BY-2:NaCl exhibited the most altered morphology (Skrzypczak et al., 2024). In BY-2:NaCl medium, concentrated NaCl impose extra threat to cells, beyond osmolarity.

Our work also demonstrates the utility of BLS and fluorescent molecular rotors in investigating viscoelasticity in plant cells cultured under various osmotic conditions. As shown, correlating Brillouin spectroscopy with fluorescence microscopy would allow for correlating mechanical properties mapping with various subcellular structures. We suppose that BLS could become a very valuable tool for revealing cell reactions that involve alterations of the cellular microenvironment.

## Discussion

Understanding the interrelations between a cellular microenvironment and an external environment remains a significant challenge in plants cell biology. One fundamental environmental factor with pleiotropic impact is osmolarity, which can rapidly alter water activity in cells, directly affect all cellular processes, and disturb biomechanical balance within plants cells elements (Watson et al., 2023; Zonia & Munnik, 2007). Plant cells sense the mechanical signals at the cell wall, plasma membrane and inside the cell. Information from these different cellular compartments is integrated to develop adaptive responses and maintain mechanical homeostasis at the cellular level (Cosgrove, 1993; Geitmann & Ortega, 2009). Although the significance of cell and tissue mechanics in plant development has been appreciated for many years (Echevin et al., 2019; Moulia et al., 2015, 2019, 2021; Trinh et al., 2021), the study of cellular micromechanics at the level of the individual cell has proven to be problematic. The universal presence of the cellulosic cell wall and the apoplastic continuity that it provides endows plant tissues with a unique level of mechanical coupling. In principle, this makes it possible for plant tissues to transmit stress-mechanical information precisely and instantaneously over multicellular distances. However, the same apoplastic continuity that makes stress-mechanical signaling attractive as a possible developmental effector also makes it difficult to interpret responses and isolate mechanical variables at the level of the individual cell. Using individual cells cultivated in suspension helps control these variables, making it easier to interpret how cells respond to a given environmental stimulus. Such model system allows to study plant biomechanical properties and their regulation by various extrinsic and intrinsic factors influenced by the physical interactions between cells and their environment (Weber et al., 2015).

A morphological and aggregation aberrances of BY-2:NaCl cells show that osmoadaptation does not necessarily result in complete reestablishing of reference cells status. In contrast to BY-2:NaCl, the BY-2:Mann morphology and growth more resembles the BY-2:Control, despite higher osmolarity of medium (Skrzypczak et al., 2024). Interestingly, in *S. cerevisiae* it was shown, that starvation leads to decreased cells size and increased molecular crowd (Joyner et al., 2016). Our results does not support correlation between small size and crowdedness of BY-2:NaCl cells. Moreover, high osmolarity of BY-2:Mann medium did not lead to the decreased cells size, so this effect might be salinity dependent disruption of cells metabolism. Does small size correspond to ‘starvation-like’ effect in BY-2:NaCl, it is still unknown.

Recent studies have demonstrated that in neuroblastoma cells hyperosmolarity rapidly increased cellular viscosity and molecular crowding (Kitamura et al., 2023). In human cell lines, the conservation of viscosity across numerous cell lines indicates the importance of maintaining viscosity within a certain range (Kwapiszewska et al., 2020). This may not be the case in plant cells, where physical forces are different and largely determined by cell walls and turgor pressure. However, the slightly altered viscoelasticity of BY-2 cultures adapted to high osmolarity highlights the potential of plant cells to maintain viscoelastic properties in the harsh environmental conditions.

The PEG-BDP rotors allowed us to discover a stable level of molecular crowding in cytoplasm of BY-2 lines. However, if consider differences between cytoplasm and environment lifetime (Δ), cytoplasm of BY-2:NaCl differed less in density from environment, than in BY-2:Control and BY-2:Mann. For all lines we observed also the dynamic responses - the increased MC after osmotic stress, by which adapted BY-2 exhibited smaller MC changes, than BY-2:Control. On the other hand, the BLS revealed that elastic contrast in the cytoplasm dominating areas was greater in adapted cells, than in BY-2:Control. However, both results suggest distinguished crowding specificity of BY-2:NaCl cytoplasm, that differ the most from BY-2:Control. The similarities and differences between data obtained with BLS and fluorescent molecular rotors require further research to point out, if they depend on various measured phenomenon, or properties of either media or cells. Relations between osmolarity, water activity, MC, viscosity, and ionic strength in cellular microenvironments are overlapping and interrelated, and challenging to elucidate *in situ*.

Vacuoles constitute a key component in plants cell osmoregulation, their content and mechanical properties are important for the stress response (Afzal et al., 2016; Maurel et al., 1997). How it could work a coordination between ions and water transport through PM and tonoplast, it has been proposed for guard cells (Cubero-Font & De Angeli, 2021). By BLS we obtained data on approximate vacuoles areas, that were distinguished from cytoplasmic bands areas pushed to cell walls by vacuole’s pressure. We did not have the possibility to integrate BLS with fluorescence microscopy, which would have allowed for a more precise identification of the subcellular compartment being measured. Both viscous and elastic contrasts are the greatest for vacuoles in BY-2:Mann. Such dense and viscous vacuoles correspond with the greatest osmolarity of BY-2:Mann medium. The high medium osmolarity impose adaptations, that would be concentrated on preventing water efflux and sustaining turgor.

We revealed interesting properties of measured by BLS elastic contrast in BY-2:NaCl. Only in this one adapted line, BLS measured molecular crowd seems to be higher in cytoplasm, than in vacuoles. It is tempting to speculate disturbed relations between biomechanics of vacuole and cytoplasm, that might lead to decreased size of BY-2:NaCl cells. However, still more specific research on relation between vacuole and cytoplasm microenvironments, is required to explain osmoadaptation and its consequences.

The adapted BY-2 lines are characterized also by different abiotic stress related gene expression patterns (Skrzypczak et al., 2024). Both adapted cell lines successfully mitigated the prolonged risk of increased molecular crowding, membrane disintegration, and other disruptions triggered by hyperosmolarity stress. It is important to note that the impacts of NaCl on cells go beyond hyperosmolarity, and includes sodium-specific toxicity and increased ionic strength, which can disturb the cellular microenvironment (Honig & Nicholls, 1995). However, at some rate mannitol also increases ionic strength by inducing water efflux from cells, followed by a regulatory volume increase through the uptake of ions, and relative increase in ionic cellular constituents (Zonia & Munnik, 2007). In future, advances in cellular biosensors of ionic strength could help to distinguish effects of ionic strength, ionic toxicity, and osmolarity/water activity on cells (Liu et al., 2017; Miller et al., 2020).

Many questions remain to be answered regarding hyperosmolarity adaptation mechanisms. For instance, how do adapted BY-2 cells maintain efficient molecular machinery despite increased media osmolarity? Is the osmosensing machinery persistently hyperactivated in adapted BY-2 cells? Although osmoregulation mechanisms in plants have yet to be fully elucidated, mechanosensitive channels likely play a crucial role in osmosensing (Gorgues et al., 2022; Mousavi et al., 2021). In animal cells, the detection of osmotic stress is associated with the activity of kinase ASK3 (MAP3K16), which depends on the location within poly(ADP-ribose) containing condensates, coupled with liquid-liquid phase separation and cascades of the signalling pathway (Naguro et al., 2012; Watanabe et al., 2021). In plants, mono- and poly(ADP-ribosyl)ation was shown as involved in immunity, that often exhibits crosstalk with abiotic stress response (Feng et al., 2015; Kong et al., 2021; Yao et al., 2021). The viscoelasticity *per se* has been indicated as an important factor in liquid-liquid phase separation processes in biological systems (Tanaka, 2022).

## Conclusions

In this paper the influence of varying environmental osmotic conditions on the long-term (adaptation) and short-term scale (stress) on the viscoelastic properties of BY-2 cells was presented and discussed. Fluorescent molecular rotors and Brillouin Light Scattering were innovatively used for analysis of viscoelasticity of living plants protoplasts with subcellular resolution. We revealed that stability of molecular crowding in cytoplasm and tension of plasma membrane correlated with long term suspension cell cultures survival and growth in environments of high osmolarities. Achieving such equilibrium in cellular microenvironment might be considered the important element of osmoadaptation. Interestingly, as a result of long-term osmoadaptation, the BY-2 lines exhibited smaller changes in cytoplasmic crowding in response to the hyperosmolarity stress. BLS analysis enables us to reveal variations in elastic and viscous contrasts between BY-2 lines. Especially, the BY-2:Mann cells are characterised by the most crowded vacuoles, which correspond with the highest media osmolarity. Many challenges remain in the field of plant mechanobiology, to which our correlative approach could provide valuable insight into plant cell responses to the environment and the further development of useful methods.

## Supporting information

SI_BDP_synthesis

SI_BLS

SI_BDPimage

## Author contributions

P.W., A.K-M., and T.S. conceptualized the study. A.K-M performed BY-2 cell adaptation. T.S. and M.P. performed the experiments with BY-2 cells and analyzed the data. M.R. synthesized BDP-molecular rotors. T.S. and M.P. wrote the original draft, and A.K-M. reviewed the manuscript.

