## Supplementary material for "The viscoelastic properties of *Nicotiana tabacum* BY-2 suspension cell lines adapted to high osmolarity": SI_BDP_synthesis

**Contents:**

1. *General information* 1

*2. Synthesis of compounds **1-7*** 2

3. *References* 6

*4. $^{1}H$ NMR, $^{19}F$ NMR and $^{13}C$ NMR spectra of compounds* 6

1. *General information*

$^{1}H$ NMR, $^{19}F$ NMR and $^{13}C$ NMR were performed on Varian VNMR-S (400 MHz), and Varian Mercury (300 MHz) spectrometers, as is noted. Chemical shifts of $^{1}H$ NMR and $^{13}C$ NMR were expressed in parts per million downfield and upfield from trace of solvent ($^{1}H$ NMR $CDCl_3$ δ 7.26, $CD_3OD$ δ 3.31) ($^{13}C$ NMR $CDCl_3$ δ 77.16, $CD_3OD$ δ 49.00) as the internal standard. Chemical shifts of $^{19}F$ NMR were expressed in parts per million. MS (ESI) spectra were performed on ZQ4000 Waters Mass Spectrometer, as well as MS(EI) on 320 MS/450 GC Bruker.

Reagent grade chemicals were used and solvents were dried by refluxing with $CaCl_2$ ($CH_2Cl_2$), with NaH (THF) and distilled under an argon atmosphere. TLC was performed on Merck Kieselgel 60-$F_{254}$ with $CH_2Cl_2$ or $CH_2Cl_2$/MeOH as developing systems, and products were detected by inspection under UV-Vis light (254 nm, 360 nm). Merck Kieselgel 60 (230–400 mesh) was used for column chromatography. All starting materials were supplied by Angene, Acros, or Merck Life Science. All moisture sensitive reactions were carried out under argon atmosphere using oven dried glassware. Compounds **1-2** [1],

**4** [2], **5** [3], **6** [2], **7** [1] were prepared as described. The $^{1}H$ and $^{13}C$ NMR data for **1**-**2**, **4-7** [1] were in good agreement.

*2. Structure of BODIPY (BDP) based molecular rotors **1-2***

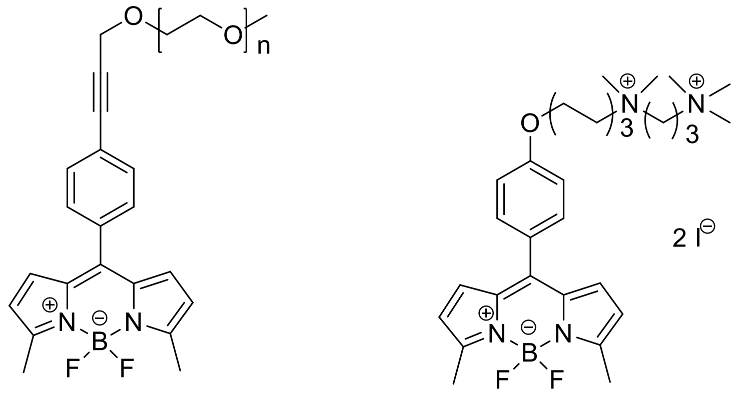

**1**
**PEG-BDP**

**2**
**N⁺-BDP**

**Figure S1.** Chemical structures of BODIPY derivatives **1**-**2** used in this study.

*2. Synthesis of BDP **5-6***

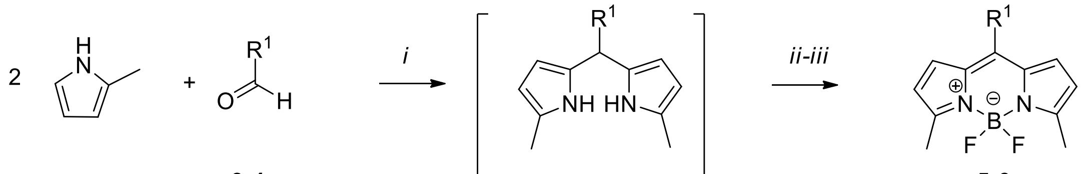

**3-4**  **5a-6a**  **5-6**

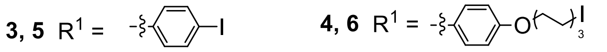

**Scheme S1**. Reaction conditions *i*. TFA, $CH_2Cl_2$; *ii*. DDQ; *iii*. (i-Pr)$_2$NEt, $BF_3$ x $Et_2O$.

*2.1. Synthesis of 5,5-difluoro-10-(4-iodophenyl)-3,7-dimethyl-5H-dipyrrolo[1,2-c:2',1'-f][1,3,2]diazaborinin-4-ium-5-uide* **5**

The solution of freshly distilled 2-methylpyrrole (210 µL, 203 mg, 2.5 mmol) and 4-iodobenzaldehyde **3** (290 mg, 1.25 mmol) in anhydrous $CH_2Cl_2$ (25 mL) was degassed by bubbling Ar for 30 min. Next, TFA (50 µL, 74 mg, 0.65 mmol) was added and a mixture was stirred for 2h. Then 2,3-dichloro-5,6-dicyano-1 4-benzoquinone (DDQ) (285 mg, 1.25 mmol) was added, the reaction was degassed by bubbling Ar for 10 min, followed by stirring for additional 20 min. Next, diisopropylethylamine (1.53 mL, 1.14 g, 8.8

mmol) was added, followed by the addition of boron trifluoride diethyl etherate (1.55 mL, 12.6 mmol), and the mixture was stirred at RT overnight. Then, the solvent was evaporated, and the residue was dissolved in $CH_2Cl_2$, purified and repurified by column chromatography ($CH_2Cl_2$) to give **5** as a deep-orange oil. (103 mg, 19%) **$^1$H NMR** (300 MHz, $CDCl_3$) δ 7.90 – 7.78 (m, 2H), 7.25 – 7.19 (m, 2H), 6.68 (d, *J* = 3.8 Hz, 2H), 6.27 (d, *J* = 4.6 Hz, 2H), 2.65 (s, 6H). **$^{19}$F NMR** (283 MHz, $CDCl_3$) δ -148.12 (q, *J* = 32.4 Hz). **$^{13}$C NMR** (101 MHz, $CDCl_3$) δ 158.04, 140.99, 137.44, 134.15, 133.52, 131.87, 130.12, 119.67, 96.35, 14.93. MS (EI) calcd. for $C_{17}H_{14}BF_2IN_2$ $[M]^+$ 422.03; found 422.1.

*2.2. Synthesis of* 5,5-difluoro-10-(4-((6-iodohexyl)oxy)phenyl)-3,7-dimethyl-5H-dipyrrolo[1,2-c:2',1'-f][1,3,2]diazaborinin-4-ium-5-uide **6**

*Step A. Synthesis of* **6a**. The solution of freshly distilled 2-methylpyrrole (0.7 mL, 676 mg, 8.33 mmol) and 4-((6-iodohexyl)oxy)benzaldehyde **4** (250 mg, 0.75 mmol) was degassed by bubbling Ar for 30 min. Next, TFA (13 µL, 19 mg, 0.17 mmol) was added and a mixture was stirred at RT for 2h and evaporated under reduced pressure at 60 °C. Then, $CH_2Cl_2$ was added, and organic layer was washed with saturated $NaHCO_3$, brine, dried ($Na_2SO_4$) and evaporated under reduced pressure. The residue was column chromatographed (*n*-hexane:$CH_2Cl_2$:1:2, v:v to give **6a** (234 mg, 65 %) as slightly brownish green oil. **$^1$H NMR** (401 MHz, $CDCl_3$) δ 7.66 (s, 2H), 7.22 – 7.13 (m, 2H), 6.93 – 6.83 (m, 2H), 5.87-5.80 (m, 2H), 5.82 – 5.75 (m, 2H), 3.98 (t, *J* = 6.4 Hz, 2H), 3.25 (t, *J* = 7.0 Hz, 2H), 2.24 (s, 6H), 1.94 – 1.80 (m, 4H), 1.55 – 1.48 (m, 4H). **$^{13}$C NMR** (101 MHz, $CDCl_3$) δ 157.94, 134.55, 131.73, 129.42, 127.08, 114.52, 107.12, 105.89, 67.81, 43.34, 33.46, 30.31, 29.16, 25.16, 13.12, 7.06.

*Step B. Synthesis of* **6**. To the solution of **6a** (157 mg, 0.3 mmol) in $CH_2Cl_2$ (8.5 mL) 2,3-dichloro-5,6-dicyano-1 4-benzoquinone (DDQ) (75 mg, 0.32 mmol) was added, the reaction was degassed by bubbling Ar for 10 min, followed by stirring for additional 50 min. Then triethylamine (138 µL, 107 mg, 1.06 mmol), was added, followed by the addition of boron trifluoride diethyl etherate (0.1 mL, 0.8 mmol), and the mixture was stirred at RT overnight. Next, the solvent was partially evaporated, and purified by column chromatography ($CH_2Cl_2$) to give **6** (82 mg, 48%) as an orange oil. **$^1$H NMR** (300 MHz, $CDCl_3$) δ 7.46 – 7.39 (m, 2H), 7.01 – 6.94 (m, 2H), 6.75 (d, *J* = 4.2 Hz, 2H), 6.26 (d, *J* = 4.2 Hz, 2H), 4.02 (t, *J* = 6.3 Hz, 2H), 3.22 (t, *J* = 6.9 Hz, 2H), 2.64 (s, 6H), 1.96 – 1.76 (m, 4H), 1.58 – 1.46 (m, 4H). **$^{19}$F NMR** (283 MHz, $CDCl_3$) δ -148.06 (q, *J* = 32.5 Hz). **$^{13}$C NMR** (101 MHz, $CDCl_3$) δ 160.89, 156.99, 142.78, 134.57, 132.13, 130.35, 126.48, 119.25, 114.30, 68.03, 33.45, 30.31, 29.08, 25.16, 14.96, 7.05. MS (EI) calcd. for $C_{23}H_{26}BF_2IN_2O$ $[M]^+$ 522.11, found 522.1.

*2.3. Synthesis of 10-(4-(2,5,8,11,14,17,20,23,26,29,32,35,38,41-tetradecaoxatetratetracont-43-yn-44-yl)phenyl)-5,5-difluoro-3,7-dimethyl-5H-dipyrrolo[1,2-c:2',1'-f][1,3,2]diazaborinin-4-ium-5-uide* **1**

*Step A. Synthesis of PEG-alkyne* **7**

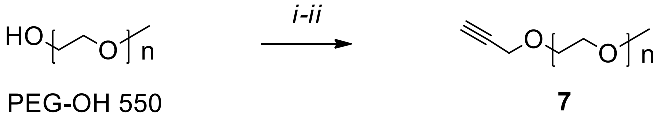


**Scheme S2.** *i*. NaH, THF, *ii*. Propargyl bromide

The mixture of NaH (60% suspension in mineral oil, 121 mg, 1.8 mmol) in anhydrous THF (20mL) was cooled down to 0°C, and degassed by bubbling Ar for 10 min. Next, PEG-OH 550g/mol (1.0 g, 1.8 mmol), was added, and a reaction mixture was stirred at 0°C for 0.5h. Then propargyl bromide (80% solution in toluene, 304 µL, 2.8 mmol) was added, and the mixture was warmed up to RT, degassed by bubbling Ar for 10 min and stirred overnight. Next, the solvent was evaporated, and the residue was dissolved in $CH_2Cl_2$. The organic phase was washed with brine, dried with $Na_2SO_4$ and evaporated under reduced pressure to get **7** (1.0 g, 92%) as un oil. **$^1$H NMR** (300 MHz, $CDCl_3$) δ 4.20 (d, *J* = 2.4 Hz, 2H), 3.74 – 3.62 (m, 52H), 3.38 (s, 3H), 2.50 (t, *J* = 2.4 Hz, 1H).

*Step B. Synthesis of **1**.*

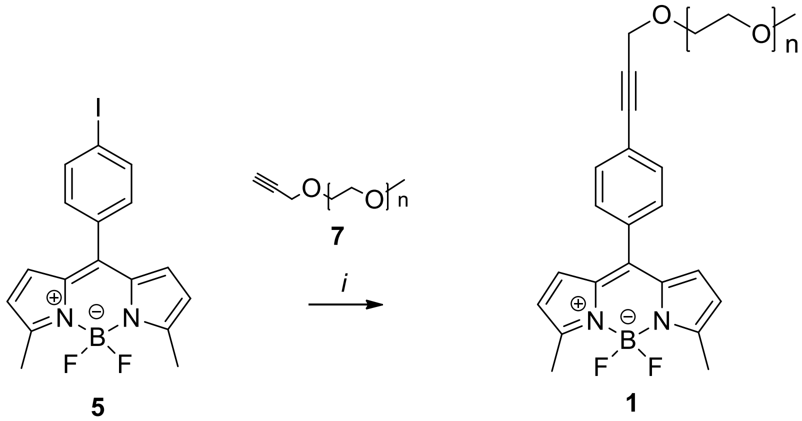


**Scheme S3.** Reaction conditions *i.* **7**, CuI, $(PPh_3)_2PdCl_2$, $(i\text{-}Pr)_2EtN$, THF.

The solution of compound **7** (155 mg, 0.26 mmol) in anhydrous THF (10 mL) was degassed by bubbling Ar for 15 min. Next, **5** (84 mg, 0.2 mmol), copper iodide (3 mg, 0.01 mmol), and bis(triphenylphosphine)palladium(II) dichloride (3 mg, 4.7 µmol) followed by diisopropylethylamine (5.7 mL, 4.21 g, 32.6 mmol) were added, and the mixture was stirred at 70 °C overnight. Then, the solvent was evaporated, and the residue was dissolved in $CH_2Cl_2$. The organic phase was washed with 1M HCl, saturated $NaHCO_3$, brine, dried ($Na_2SO_4$) and evaporated under reduced pressure. Column chromatography ($CH_2Cl_2$→$CH_2Cl_2$ : MeOH, 95:5, v:v) gave **1** (144 mg, 81%) as un orange-red oil. **$^1$H NMR** (300 MHz, $CDCl_3$) δ 7.55 – 7.49 (m, 2H), 7.44 – 7.39 (m, 2H), 6.66 (d, *J* = 4.2 Hz, 2H), 6.24 (d, *J* = 4.2 Hz, 2H), 4.44 (s, 2H), 3.72 – 3.51 (m, 52H), 3.35 (s, 3H), 2.62 (s, 6H). **$^{19}$F NMR** (283 MHz, $CDCl_3$) δ -148.12 (q, *J* = 32.4 Hz). **$^{13}$C NMR** (101 MHz, $CDCl_3$) δ 158.04, 141.62, 134.41, 134.24, 131.64, 130.46, 124.67,

119.72 (d, $J$ = 4.4 Hz), 87.46, 85.61, 71.99, 70.46-70.61 (m), 69.42, 59.31, 15.05. MS (EI) calcd for $C_{47}H_{71}BFN_2O_{14}$ $[M-F]^+$ 917.4 found 917.3.

*2.4. Synthesis of mono(10-(4-((6-(dimethyl(3-(trimethylammonio)propan-1-id-1-yl)ammonio)hexyl)oxy)phenyl)-5,5-difluoro-3,7-dimethyl-5H-dipyrrolo[1,2c:2',1'f][1,3,2]diazaborinin--4-ium-5-uide) diiodide* **2**

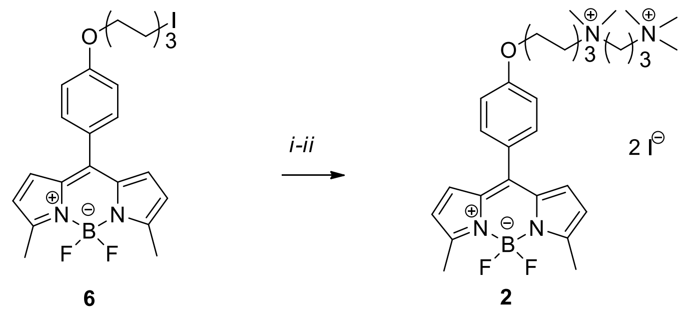


**6** **2** **N⁺-BDP**

**Scheme S4.** Reaction conditions *i.* $(CH_3)_2N-(CH_2)_3-N(CH_3)_2$, THF, *ii.* MeI, DMF.

To the solution of **6** (80 mg, 0.16 mmol) in freshly distilled THF (8 mL) tetramethylpropanediamine (2 mL) was added. The mixture was stirred at room temperature for 20h. Next, the solvent and excess of amine were evaporated under reduced pressure. The residue was washed with diethyl ether (3 x 5 mL), and dried. Then, dimethylformamide (2 mL) and iodomethane (1 mL, 2.28 g, 16 mmol) were added and the reaction was stirred at RT overnight. The solvent and excess of MeI were evaporated under reduced pressure and the residue was column chromatographed on silica gel (methanol) followed by RP C-18 silica gel ($H_2O$ → $CH_3CN$: $H_2O$, 2:8, v:v) and evaporated to get **2** as a deep red solid (57 mg, 47% yield). **$^1$H NMR** (300 MHz, $CD_3OD$) δ 7.51 – 7.46 (m, 2H), 7.11 – 7.03 (m, 2H), 6.80 (d, $J$ = 4.3 Hz, 2H), 6.36 (d, $J$ = 4.2 Hz, 2H), 4.11 (t, $J$ = 6.2 Hz, 2H), 3.61 – 3.45 (m, 6H), 3.27 (s, 9H), 3.21 (s, 6H), 2.58 (s, 6H), 2.48-2.38 (m, 2H), 1.98 – 1.85 (m, 4H), 1.72 – 1.63 (m, 2H), 1.62 – 1.47 (m, 2H). **$^{19}$F NMR** (283 MHz, $CD_3OD$) δ -146.20 (q, $J$ = 31.7 Hz). **$^{13}$C NMR** (76 MHz, $CD_3OD$) δ 162.43, 157.89, 144.22, 135.41, 133.25, 131.47, 127.13, 120.31 (d, $J$ = 3.4 Hz), 115.58, 69.12, 66.16, 63.68, 61.22, 54.51, 52.05, 29.88, 26.89, 26.56, 23.67, 19.49, 14.86. MS(ESI) calcd. for $C_{31}H_{47}BF_2IN_4O^+$ $[M-I]^+$ 667.3 found 667.

*3. References*

[1] Michels, L., Gorelova, V., Harnvanichvech, Y., Borst, J. W., Albada, B., Weijers, D., Sprakel, J. Complete microviscosity maps of living plant cells and tissues with a toolbox of targeting mechanoprobes. *Proc. Natl. Acad. Sci. U.S.A.,* **2020**, 117(30), 18110-18118. doi:10.1073/pnas.1921374117.

[2] López-Duarte, I., Vu, T. T., Izquierdo, M. A., Bull, J. A. Kuimova, M. K. A molecular rotor for measuring viscosity in plasma membranes of live cells. *Chem. Commun*. **2014**, 50, 5282-5284. doi:10.1039/C3CC47530A.

[3] Wagner, R.W.; Lindsey, J.S. Boron-Dipyrromethene Dyes for Incorporation in Synthetic Multi-Pigment Light-Harvesting Arrays. *Pure Appl. Chem.* **1996**, *68*, 1373–1380, doi:10.1351/pac199668071373.

*4. $^{1}$H NMR, $^{19}$F NMR and $^{13}$C NMR Spectra of Compounds*

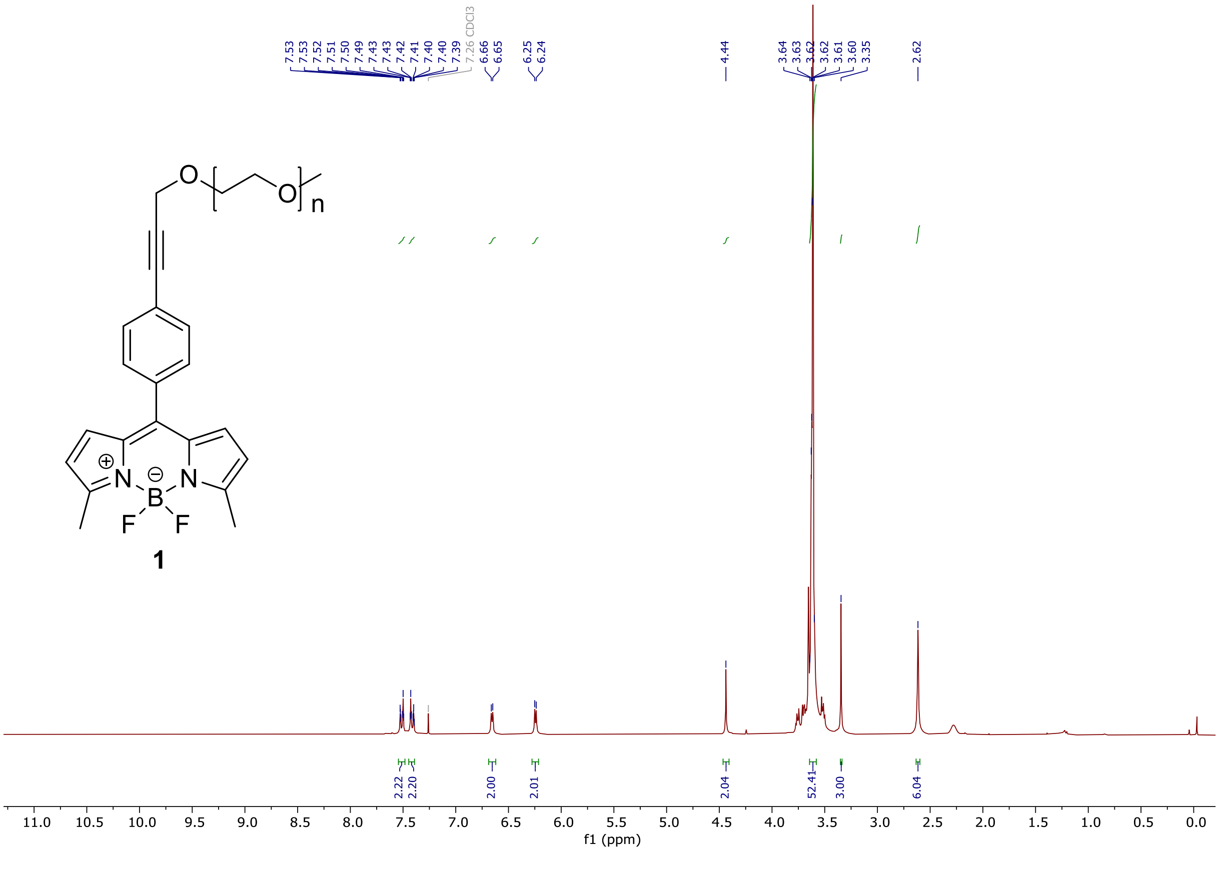


$^{1}$H NMR spectra of **1**

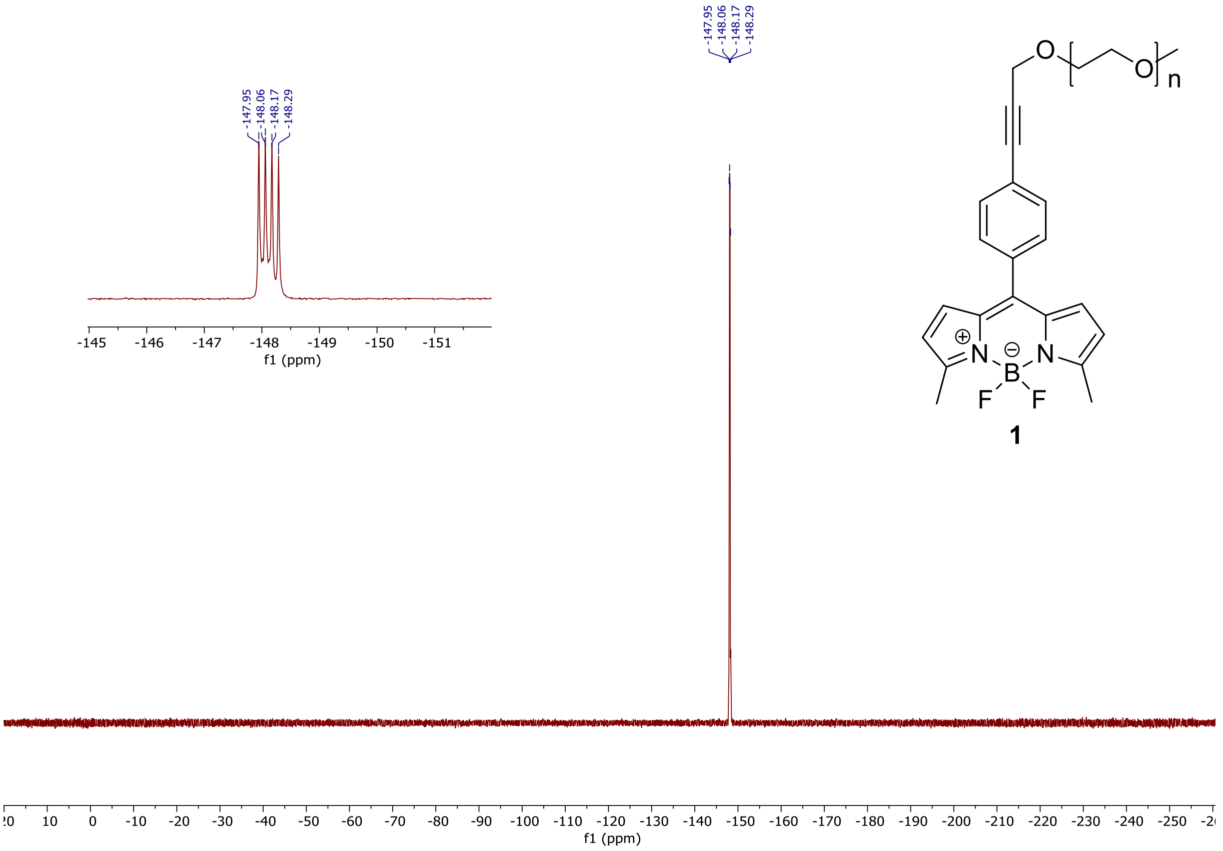


$^{19}$F NMR of **1**

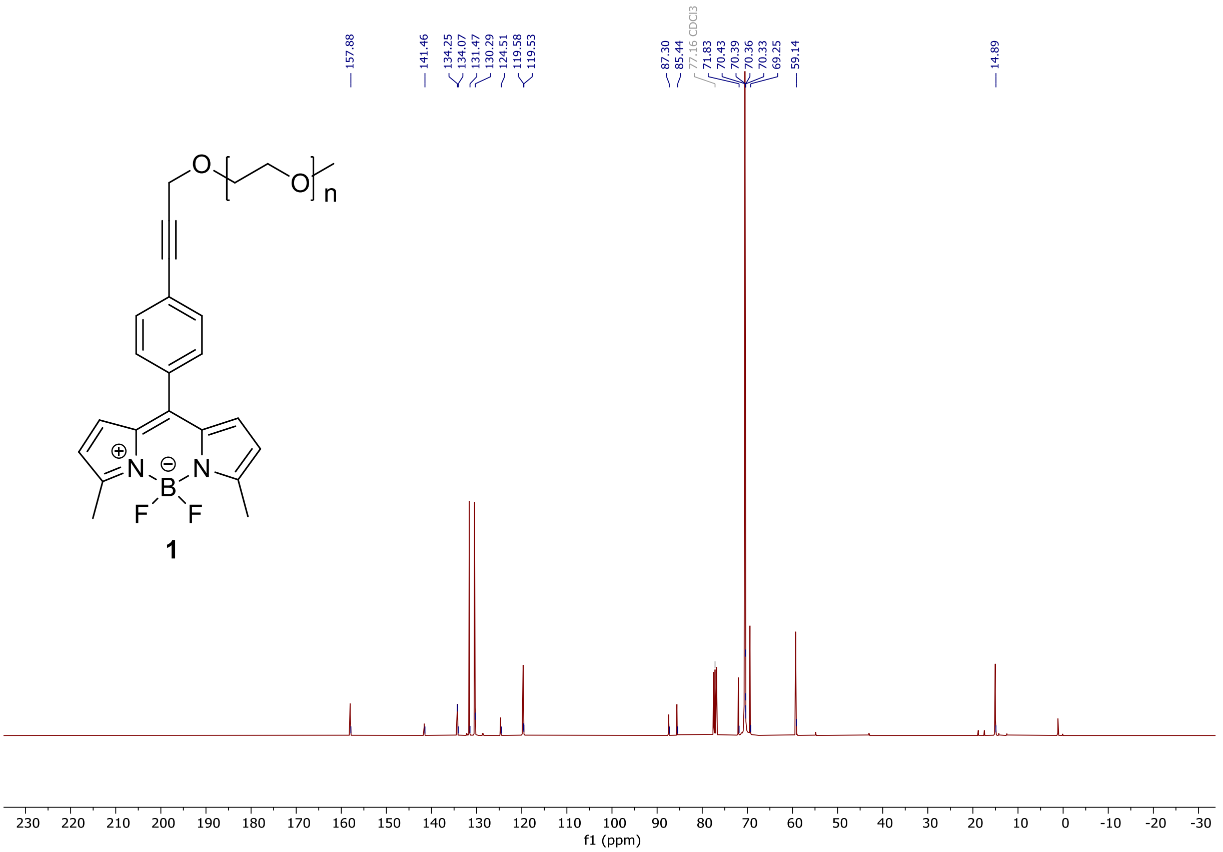


$^{13}$C NMR spectra of **1**

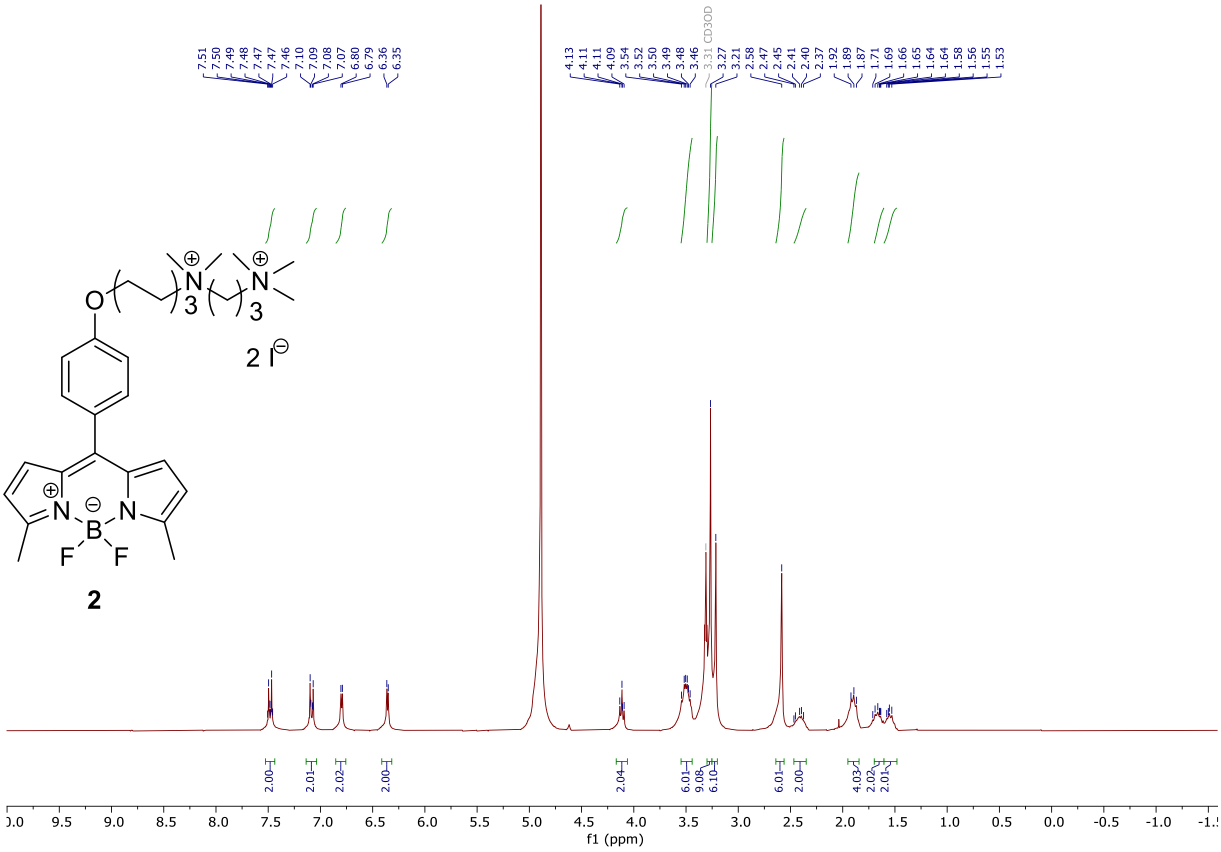


$^1$H NMR spectra of **2**

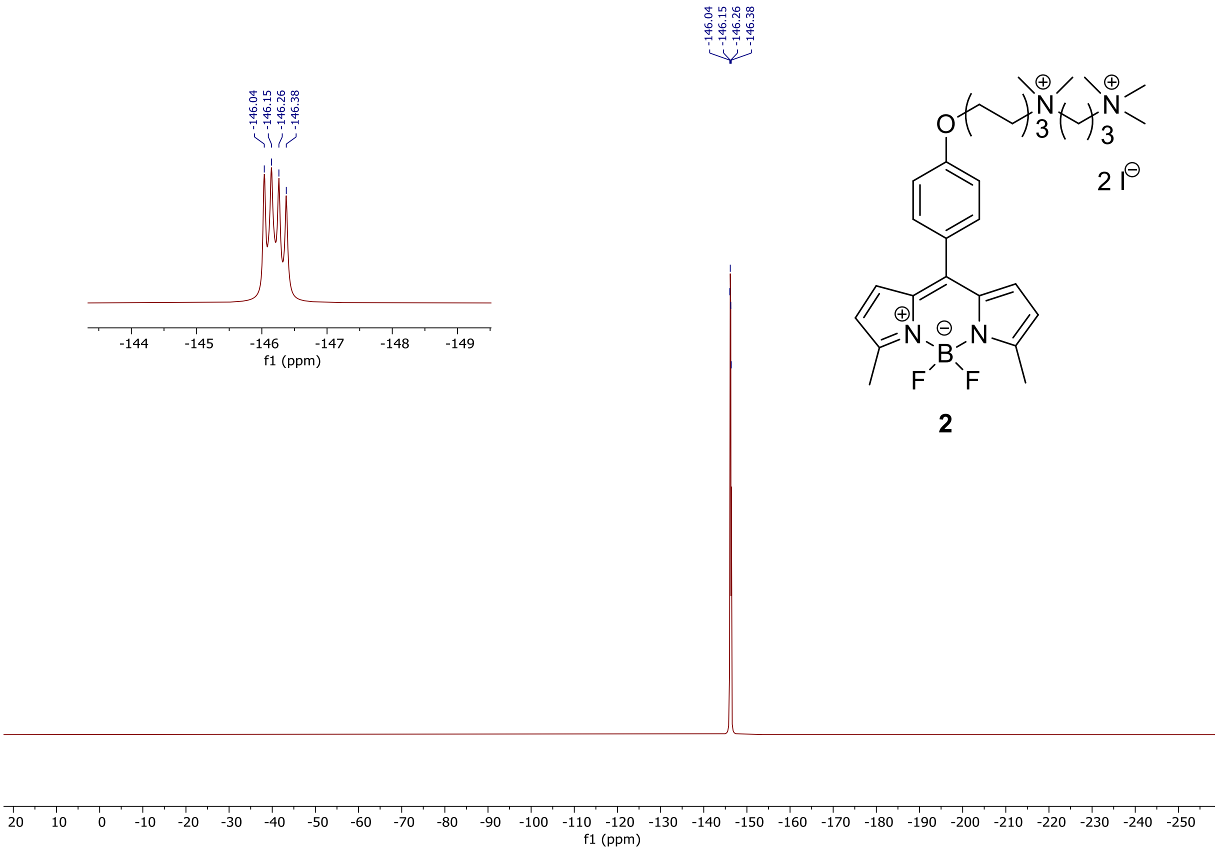


$^{19}$F NMR spectra of **2**

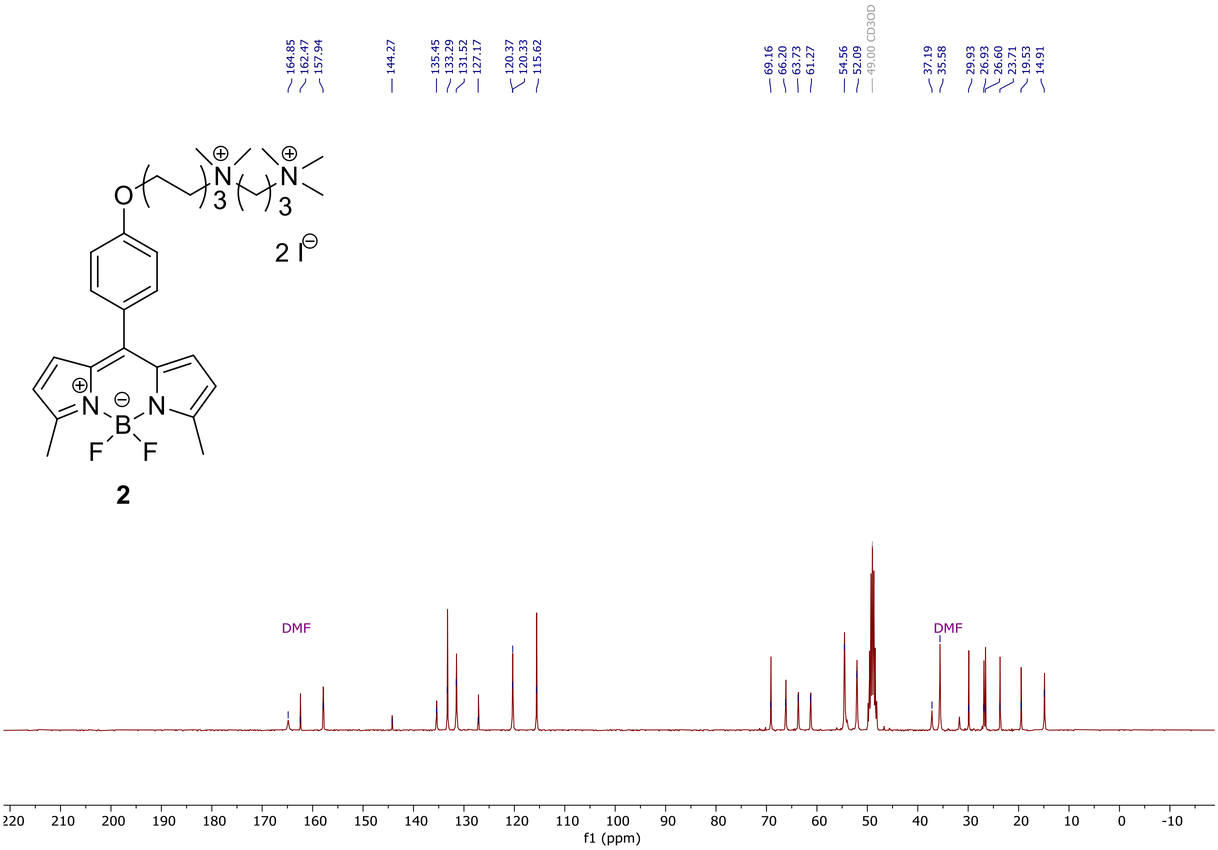


$^{13}$C NMR spectra of **2**

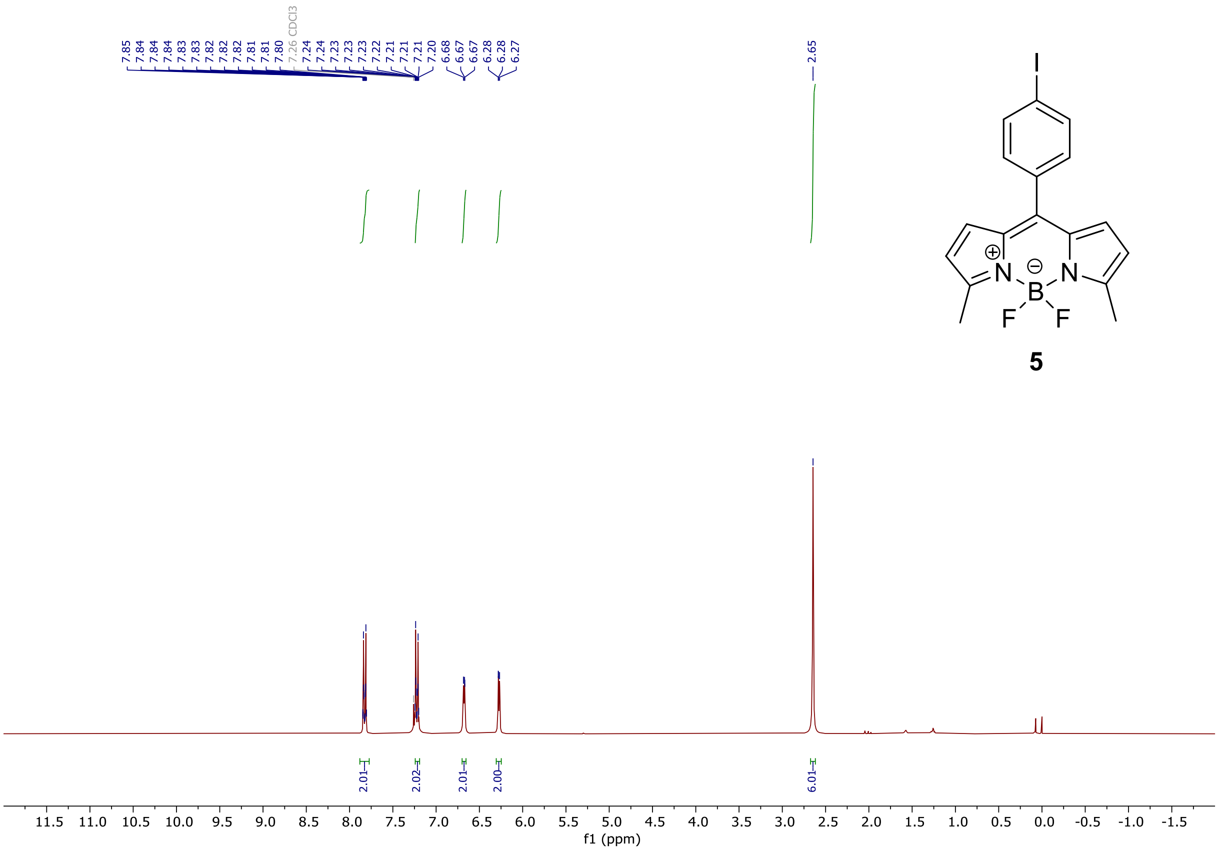


$^{1}$H NMR spectra of **5**

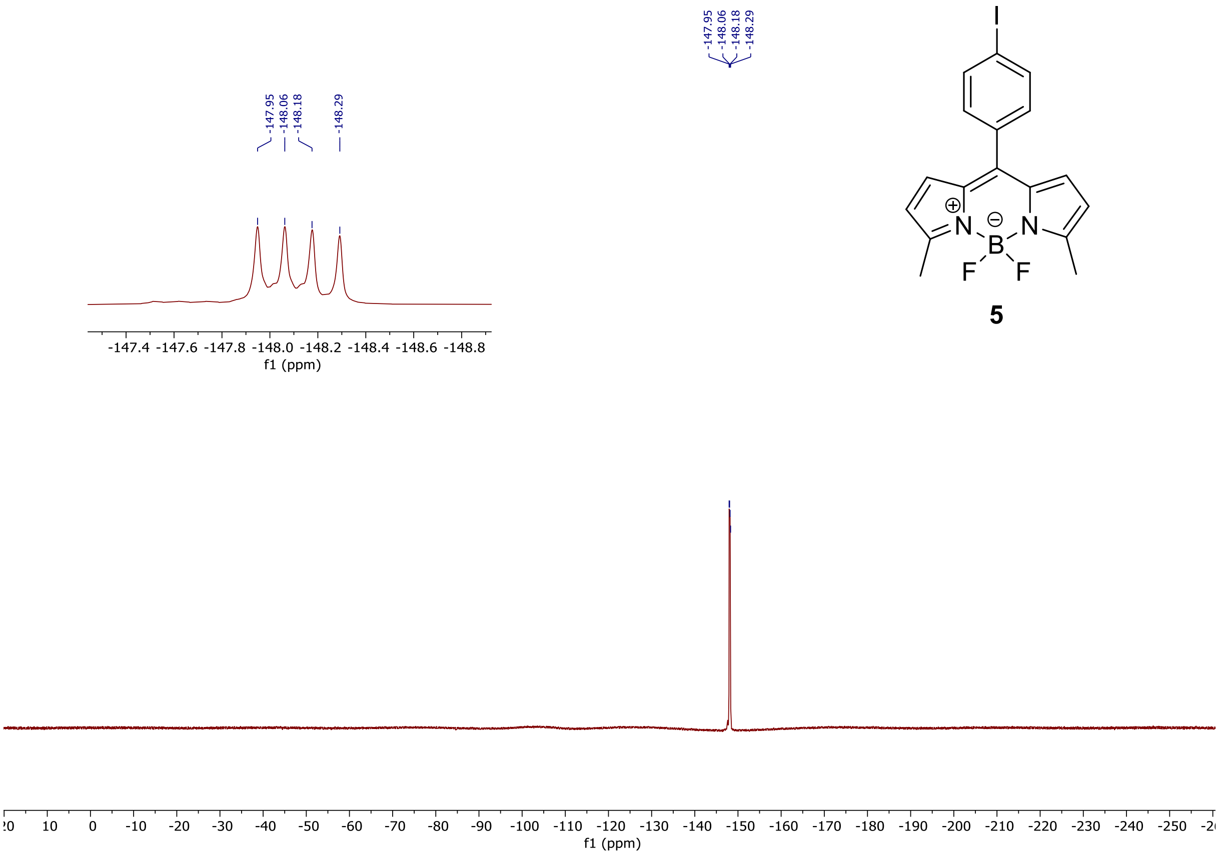


$^{19}$F NMR spectra of **5**

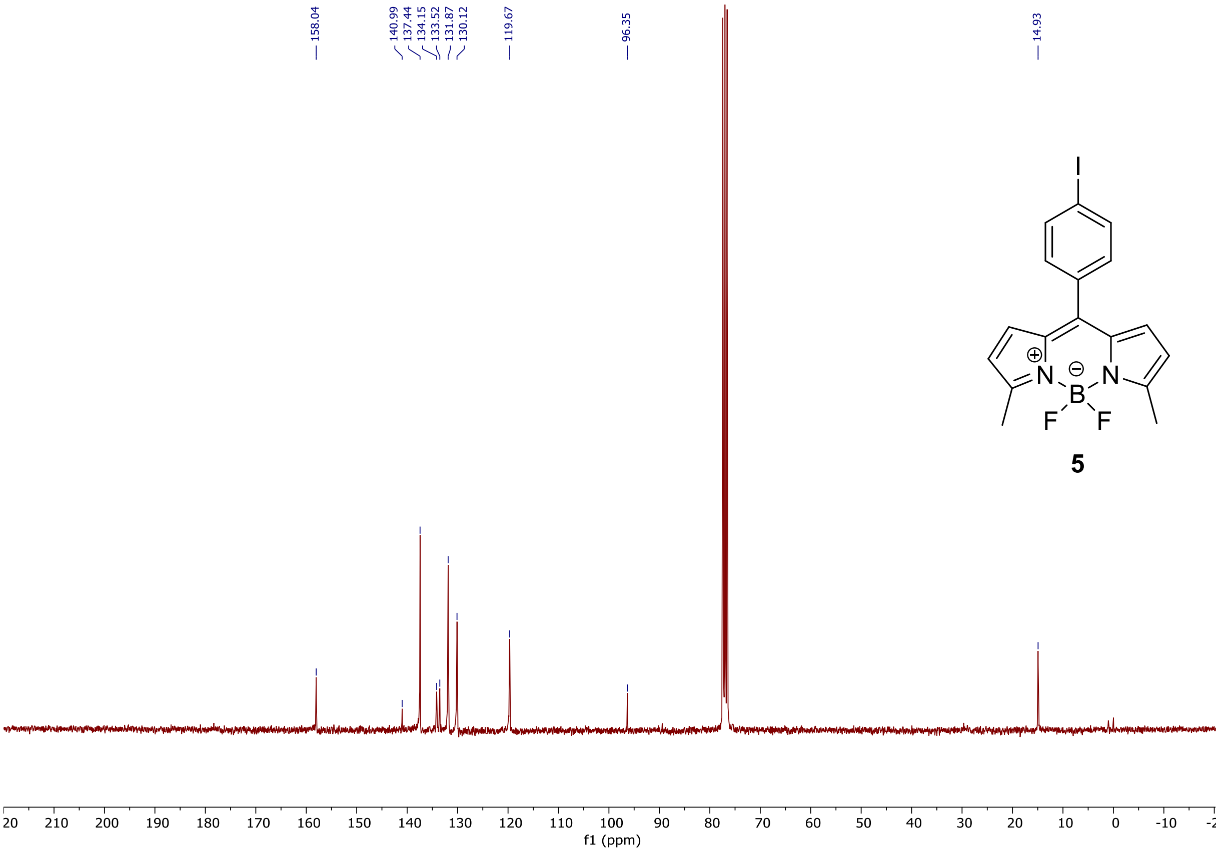


$^{13}$C NMR spectra of **5**

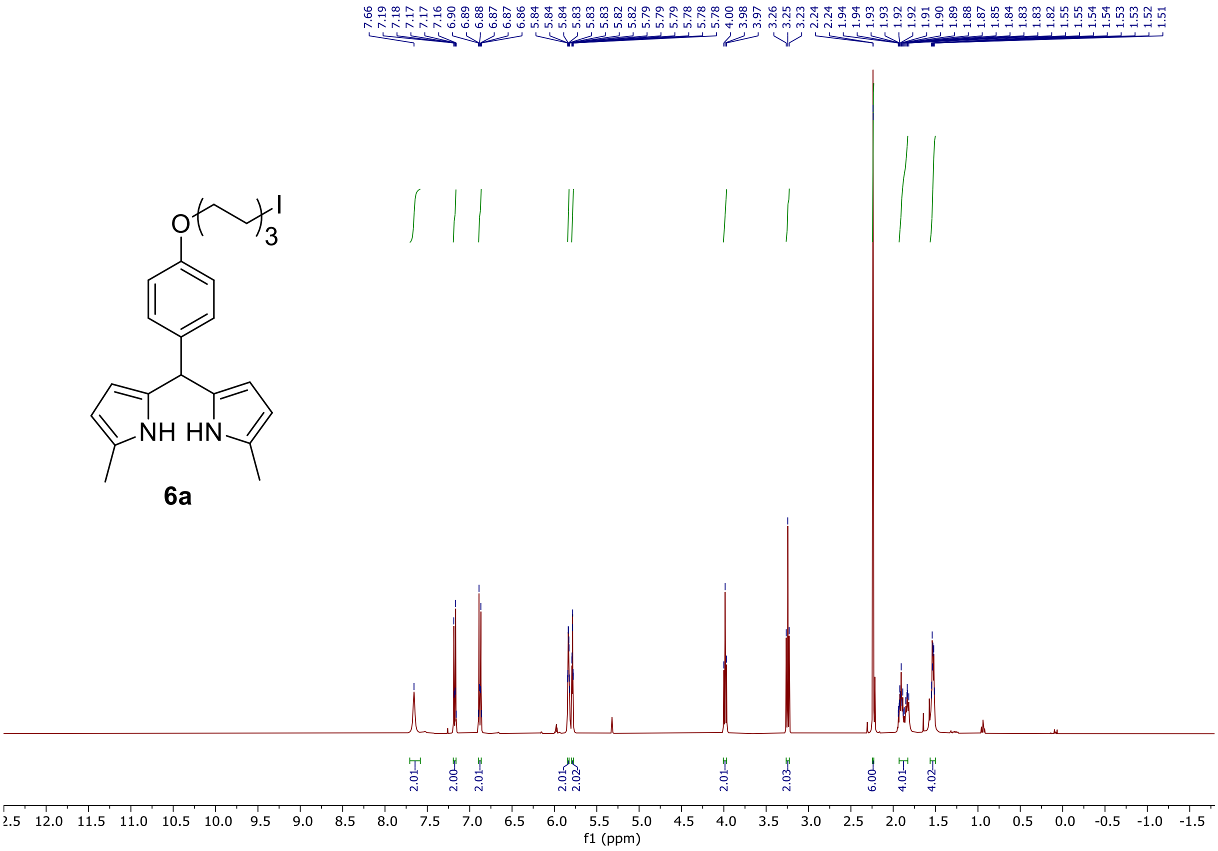

$^{1}$H NMR spectra of **6a**

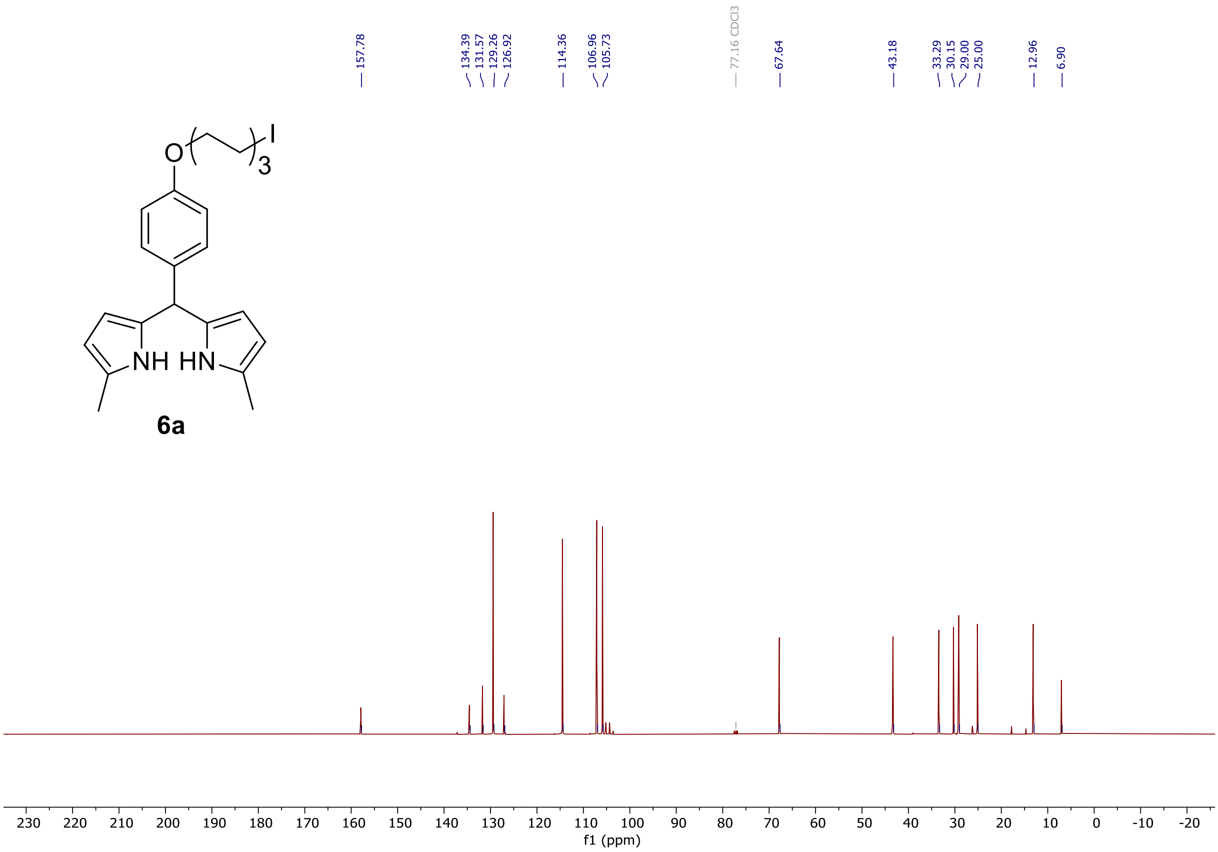

$^{13}$C NMR spectra of **6a**

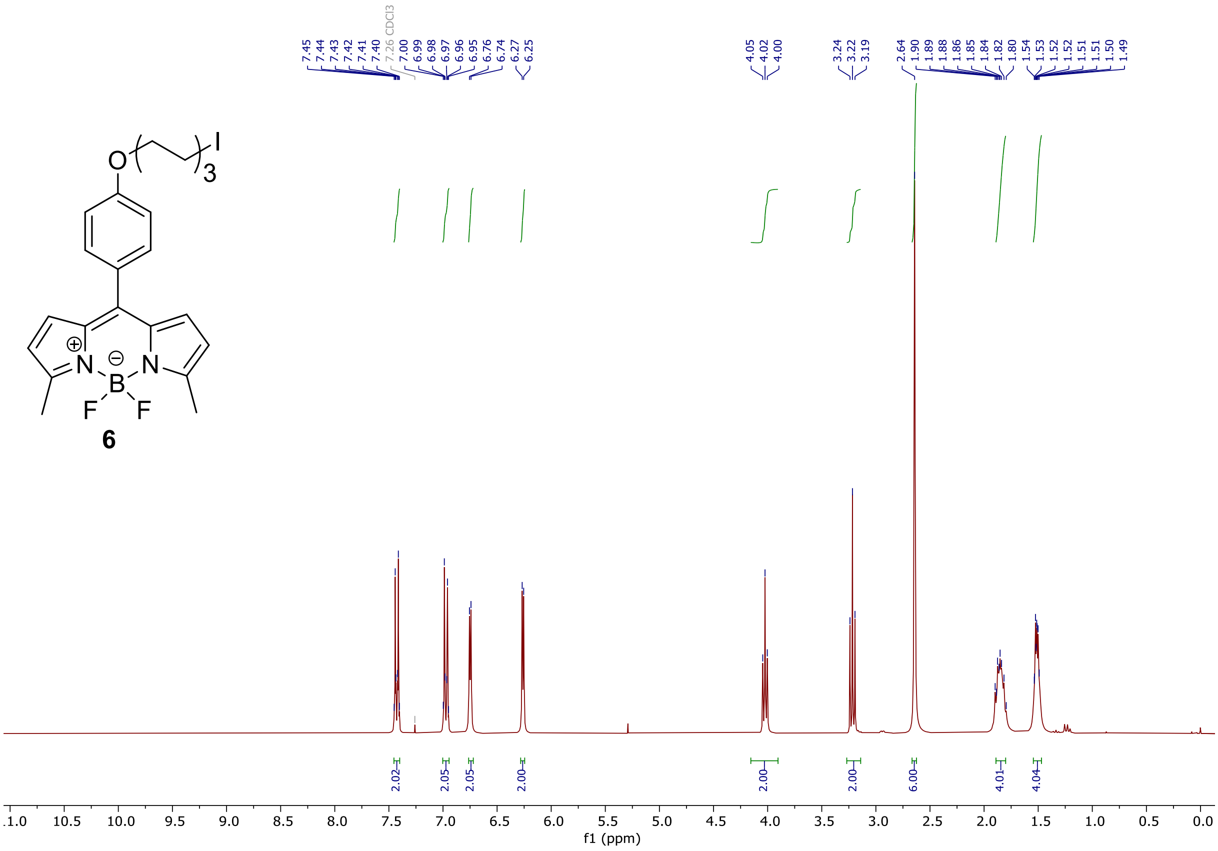


$^{1}$H NMR spectra of **6**

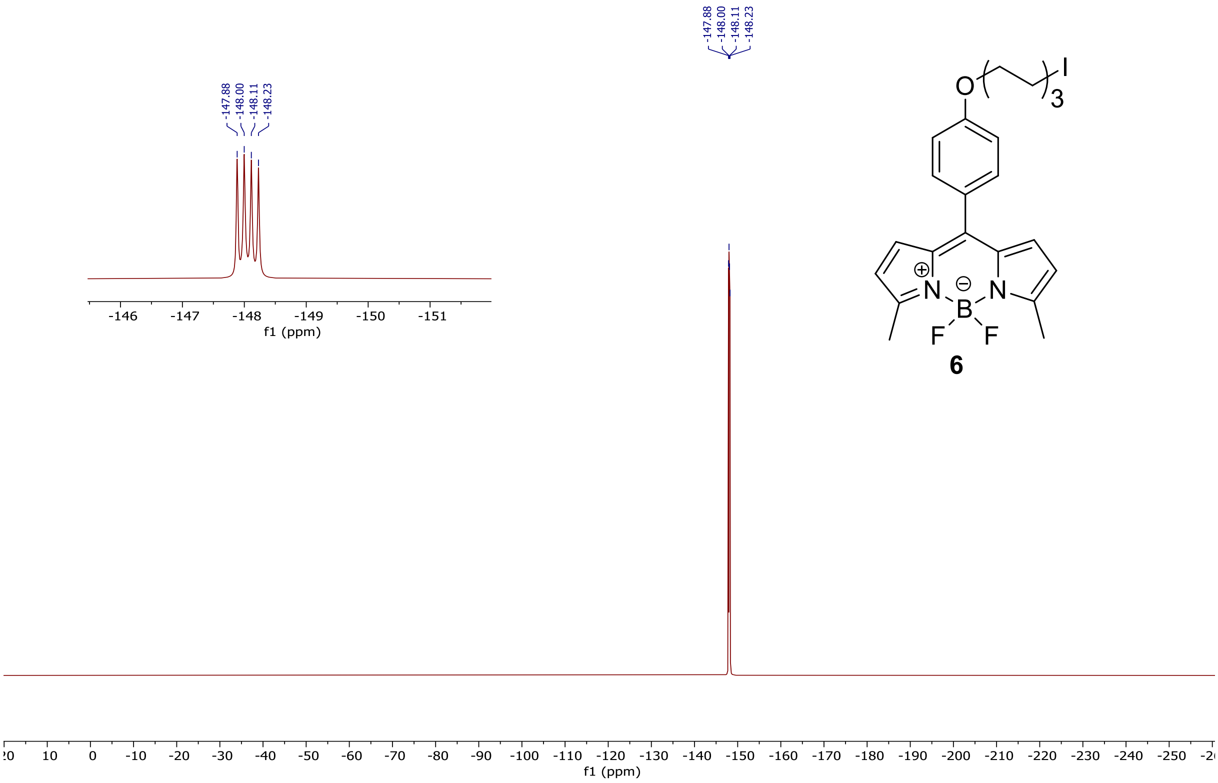


$^{19}$F NMR spectra of **6**

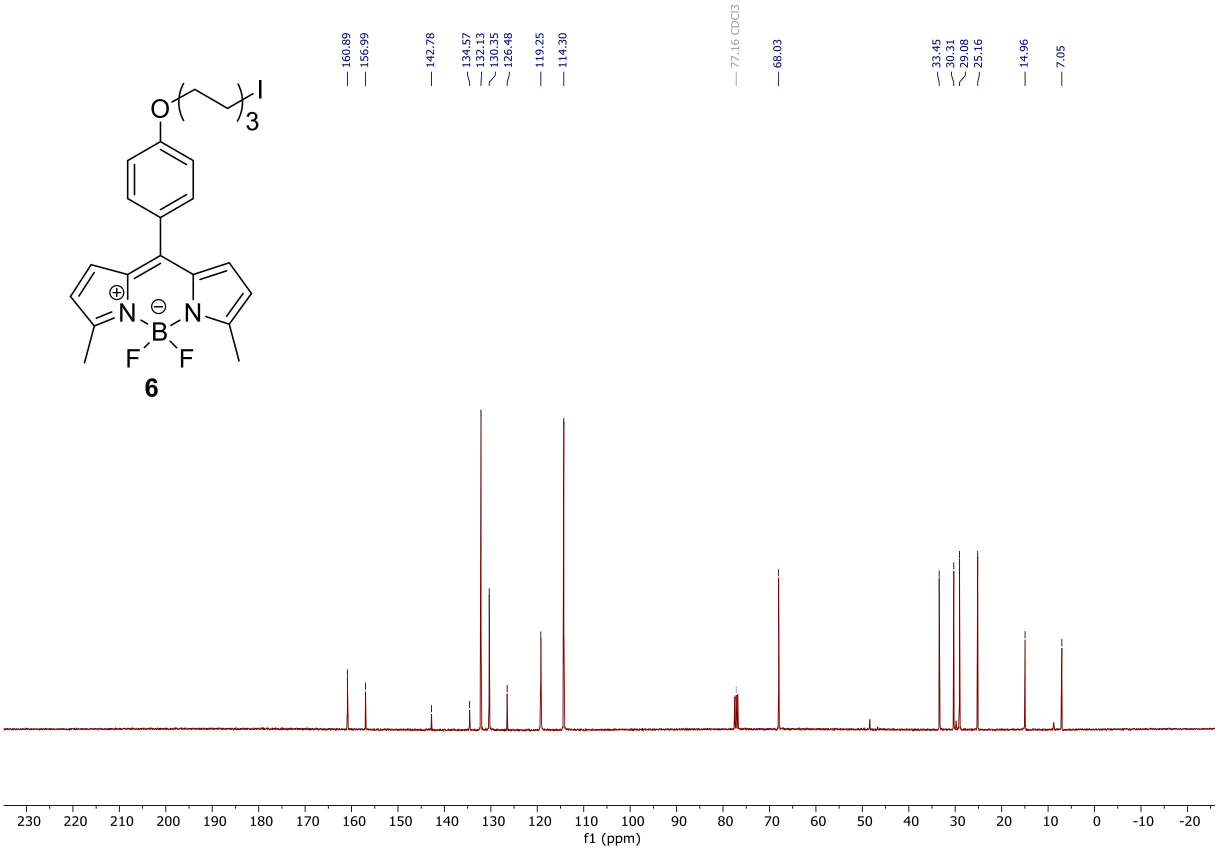


$^{13}$C NMR spectra of **6**

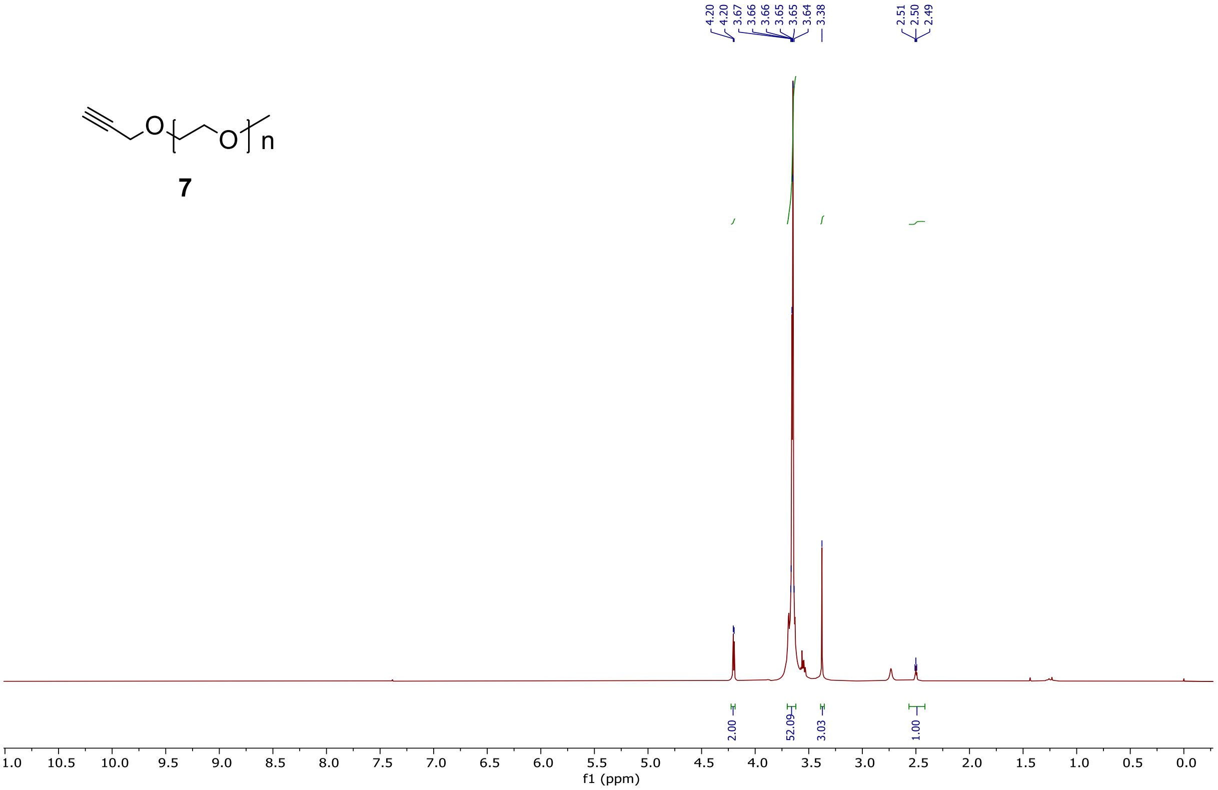


$^{1}$H NMR spectra of **7**
