## Supplementary material for "The viscoelastic properties of *Nicotiana tabacum* BY-2 suspension cell lines adapted to high osmolarity": SI_BLS

**Brillouin spectra fitting**

The Brillouin spectrum recorded from pure medium (Fig.1c) shows a single Brillouin line. To obtain the frequency shifts and linewidth, the spectra were fitted using hydrodynamic expression convoluted with the instrumental resolution function [Pecora].

$$I(f) = A_B\left[\frac{g_B}{(f+f_B)^2+g_B^2}+\frac{g_B}{(f-f_B)^2+g_B^2}\right]+$$
$$\frac{A_B g_B}{f_B}\left[\frac{f+f_B}{(f+f_B)^2+g_B^2}-\frac{f-f_B}{(f-f_B)^2+g_B^2}\right] \quad \text{(S1)}$$

In eq.S1 first two terms describe the Brillouin doublet composed of two lines symmetrically of amplitude $A_B$ shifted at $f_B$, with the width (half width at half maximum) $g_B$, whereas the last two terms ensure the preservation of the first moment sum rule and only affect the symmetry of the Brillouin doublet.

When the Brillouin signal was acquired from within the cell (Fig.1d), we found that Eq.S1 does not provide a proper description. When observed in logarithmic scale, the spectrum is clearly composed of two peaks. The spectra were fitted with the two-phase model (a sum of two Brillouin doublets) convolved with the instrumental resolution function of TFPI.

$$I(f) = \sum_{i=buffer,\mathrm{cell}}\left\{A_{B,i}\left[\frac{g_{B,i}}{(f+f_{B,i})^2+g_{B,i}^2}+\frac{g_{B,i}}{(f-f_{B,iB})^2+g_{B,i}^2}\right]+\right.$$
$$\left.\frac{A_{B,i}g_{B,i}}{f_{B,i}}\left[\frac{f+f_{B,iB}}{(f+f_{B,i})^2+g_{B,i}^2}-\frac{f-f_{B,i}}{(f-f_{B,i})^2+g_{B,i}^2}\right]\right\} \quad \text{(S2)}$$

The meaning of the symbols in Eq.S2 is the same as for Eq.S1.

We found that the parameters ($f_B$ and $g_B$) of one of these peaks correspond to those found for pure medium. Therefore, we interpret the spectra as being recorded from a two-phase system. One phase corresponds to the medium surrounding cell and the other to the cell itself. The reason we see the signal from both phases (buffer and cell) is low optical resolution of our system given by a low numerical aperture of the objective together with a high diameter of entrance pinhole (200um).

The existence of a Brillouin signal from the cell and from the buffer was previously observed in microspectroscopic experiments utilising TFPI (Mattana et al., 2018). It is known to deliver spectra of exceptional quality and it is not clear if the same “buffer contamination” exists also in spectra recorded with VIPA spectrometers more commonly utilised by Brillouin imaging.

During fitting, the Brillouin line shape parameters for the buffer phase ($f_{B,\mathrm{buffer}}$ and $g_{B,\mathrm{buffer}}$) were kept constant and fixed to the value found from the fitting spectra acquired in pure media using eq.S1.

The parameters for the first Brillouin doublet were constrained to values obtained for the medium ($f_{B,\mathrm{buffer}}$ and $g_{B,\mathrm{buffer}}$). The fitting process involved free parameters, including the second phase parameters ($f_{B,\mathrm{cell}}$ and $g_{B,\mathrm{cell}}$) and the amplitude of the Brillouin peak of the medium ($A_{B,\mathrm{buffer}}$), following a similar approach presented in a prior study (Mattana et al., 2018).

**Estimation of cells elastic and vicious contrasts**

The values of both Brillouin line parameters ($f_B$ and $g_B$) measured inside the cells (belonging to different adaptation lines) are distinctly higher than those found in their respective pure media buffers (Fig.5). At the same time, these parameters measured for bulk media were also different.

To get meaningful information on the visco-elastic behaviour of the cells adapted to mechanically different environments, the Brillouin results were presented in the form of the relative (with respect to buffer) changes of Brillouin shift and Brillouin line width

$$\upsilon_B = f_B / f_{\mathrm{B},buffer} - 1$$
$$\gamma_B = g_{\mathrm{B}} / g_{\mathrm{B},buffer} - 1 \quad \text{(S3)}$$

We call such defined properties 'the elastic contrast' $\upsilon_B$, and “the viscous contrast”, $\gamma_B$, (Bacete et al., 2021)).

It should be emphasised that the definitions of mechanical contrasts (elastic and viscous) adopted in this study differ from the usual ones (Antonacci et al., 2020), where changes in Brillouin line shape parameters are given in relation to the values characterising pure bulk water.

We decided to express the contrasts relative to the media, as these are the natural environments of the examined cells. Such defined mechanical contrasts can be interpreted in terms of change of internal cells composition being the result of cells adaptation to different environments.
