## Supplementary material for "The viscoelastic properties of *Nicotiana tabacum* BY-2 suspension cell lines adapted to high osmolarity": SI_BDPimage

1) **The relative to medium BDP-rotor lifetime**

Δ presented in percents enabled us to show cytoplasmic crowding in context of simultaneously measured media properties. PEG-BDP was shown to efficiently work in broad range of viscosities within tested water-glycerol mixtures (Michels et al., 2020).

$$\text{Delta } (\Delta) = \frac{cytoplasm\ mean\ lifetime}{medium\ mean\ lifetime}$$

2) **PEG-BDP lifetime distributions of BY-2 cytoplasm and surrounding cells media in stress conditions**

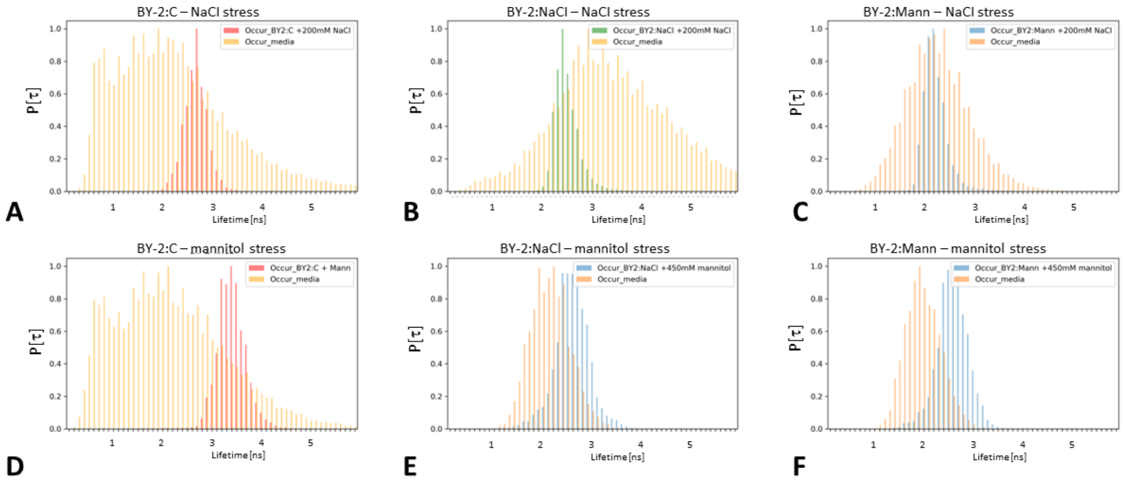


*Fig. 1. Normalised lifetime distributions of the PEG-BDP signal derived from the cytoplasm of BY-2 lines that were subjected to 200 mM NaCl stress (A-C) and 450 mM mannitol stress (D-F). The cytoplasmic PEG-BDP lifetime signals distributions are plotted on the background of the media signal derived from the vicinity of the cells (orange).*

3) **The distributions of plasma membrane $N^{+}$-BDP lifetime values during mannitol stress in BY-2:Control and BY-2:NaCl**

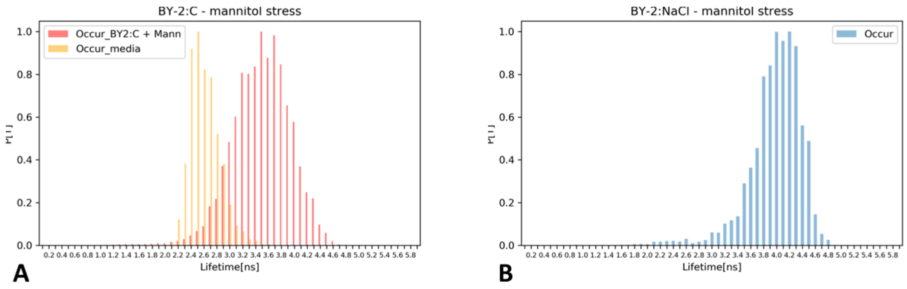


*Fig. 2. $N^{+}$-BDP lifetime histograms of BY-2:C (A) and BY-2:NaCl (B) during 450 mM mannitol stress.*
